# Loss of the UFM-1 pathway enhances the ER Unfolded Protein Response under stress to improve proteostasis and organismal health in *Caenorhabditis elegans*

**DOI:** 10.64898/2026.10.16.759640

**Authors:** Samantha Moores, Betty Ortiz-Gonzalez, Peter Juo

## Abstract

The ubiquitin-like molecule (UBL) Ubiquitin-Fold Modifier 1 (UFM1) has emerged as an important regulator of protein quality control at the endoplasmic reticulum (ER). In humans, dysfunction of the UFM1 pathway is associated with neurodevelopmental diseases and cancer, however the physiological functions of the UFM-1 pathway are not well understood. Here, we used loss-of-function mutants in *C. elegans* to investigate the role of the UFM-1 pathway in protein homeostasis and organismal health. We found that UFM-1 pathway mutants exhibit enhanced activation of the ER Unfolded Protein Response (UPR^ER^) in the presence of proteotoxic stress. Loss of the UFM-1 pathway resulted in increased numbers of polyglutamine (polyQ) aggregates in muscle and a change in aggregate composition, consistent with the regulation of protein homeostasis. Interestingly, loss of the ER-anchored components of the UFM-1 pathway also increased polyQ aggregate number, suggesting that the pathway acts at the ER to regulate cytoplasmic protein aggregates. These effects are mediated by the ribosomal subunit RPL-26, a major UFMylation target involved in ribosome-associated quality control. Tissue-specific rescue experiments indicate that the UFM-1 pathway can act cell autonomously and non-cell autonomously from neurons to regulate polyQ aggregates in muscle. Physiologically, UFM-1 pathway mutants exhibit increased survival under proteotoxic stress and increased intestinal barrier integrity in aged animals. Intriguingly, loss of the UFM-1 pathway counteracts the detrimental effects of polyQ expression on lifespan. Our study suggests that under conditions of proteotoxic stress, inhibition of the UFM-1 pathway enhances activation of the UPR^ER^ and increases animal healthspan and lifespan.

**Article summary:** The accumulation of damaged proteins is a hallmark of aging and neurodegenerative disorders such as Huntington’s disease. In this study, we tested how loss of UFMylation, a post-translational modification involved in protein quality control, influences cellular responses to proteotoxic stress, protein aggregation and aging in the nematode *C. elegans*. We found that animals lacking genes required for UFMylation had an increased number of protein aggregates and altered composition. These UFMylation mutants also had healthier aging and longer lifespan, suggesting that the UFMylation pathway is an important protein quality control mechanism that impacts organismal health and aging.

**Graphical Abstract:** 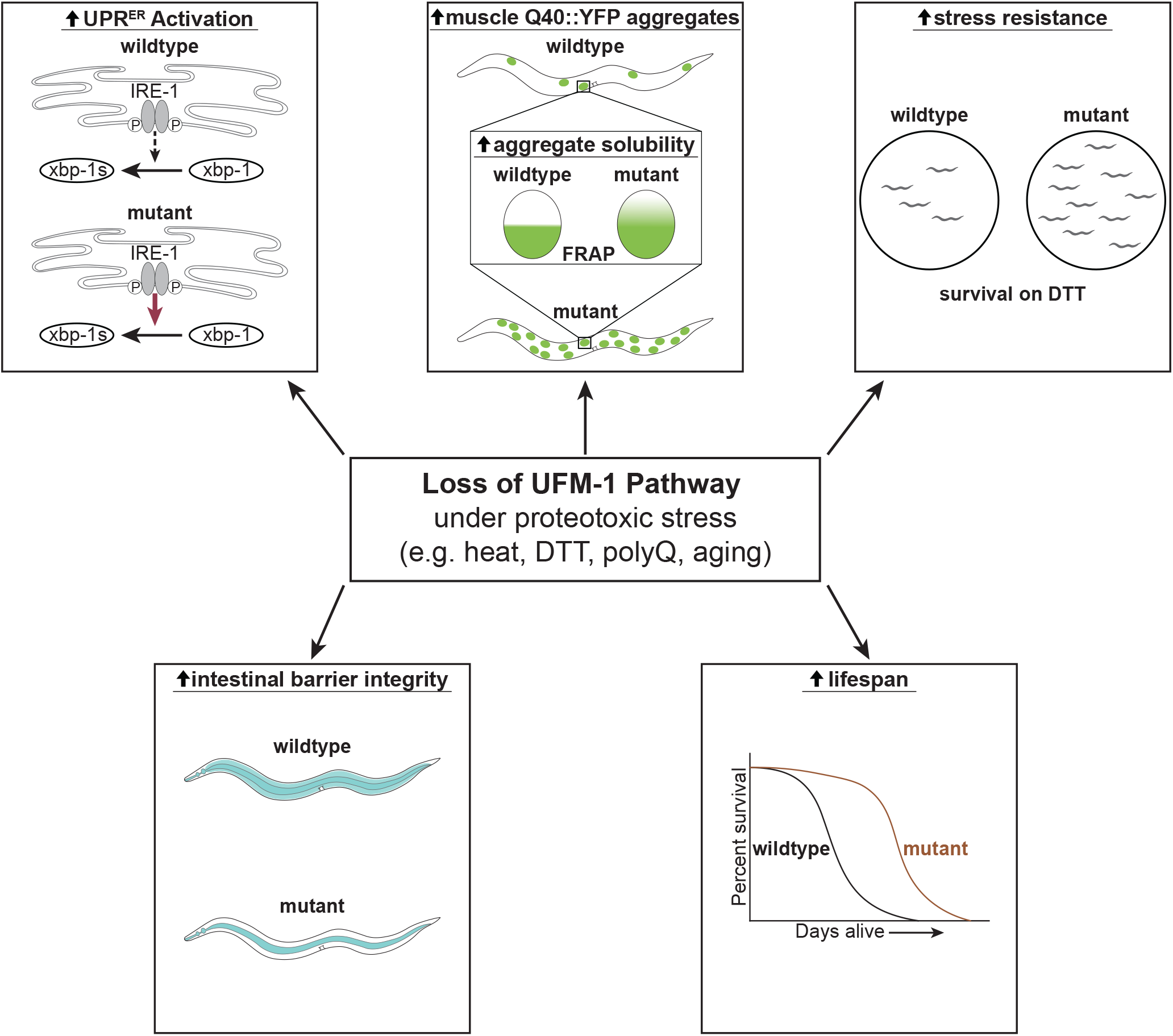

## Introduction

Dynamic regulation of the proteome, called protein homeostasis or proteostasis, is critical for cell health and viability. The proteostasis network (PN) maintains the functionality of the proteome by tightly regulating quality control mechanisms to coordinate processes ranging from protein synthesis and turnover to monitoring and regulating the accumulation of misfolded or damaged proteins (Powers and Balch 2013; Morimoto 2020; Khan 2026). When quality control and cellular degradation mechanisms are overwhelmed, misfolded or damaged proteins can form aggregates that lead to proteotoxicity and cell death (Morimoto 2020; Hetz and Dillin 2025; Khan 2026). The decline in the capacity of the PN to deal with misfolded or damaged proteins is a hallmark of aging, and results in the pathological accumulation of protein aggregates observed in many age-related neurodegenerative disorders such as Huntington’s Disease (HD) and Alzheimer’s Disease (Taylor and Hetz 2020; López-Otín et al. 2023; Wilson et al. 2023; Khan 2026).

Polyglutamine (PolyQ) expansion diseases, like HD, are caused by CAG repeats which encode long glutamine tracts that are prone to protein misfolding and aggregation (Andrew et al. 1993; Tabrizi et al. 2020; Johnson et al. 2022). PolyQ aggregation increases with age and disease severity and is dependent on the number of repeats with ∼35-40 CAG repeats typically considered the disease threshold for HD (Andrew et al. 1993; Morley et al. 2002; Johnson et al. 2022). PolyQ aggregation models have been established in *C. elegans* where fluorescently-tagged polyQ proteins with various numbers of CAG repeats can be expressed and monitored *in vivo*. Similar to the human diseases, polyQ aggregation severity in these models is dependent on age and the number of CAG repeats. These models have been used to identify genes and mechanisms that regulate polyQ aggregation in *C. elegans*, including those that act both cell-autonomously and non-cell autonomously (Garcia et al. 2007; Taylor and Dillin 2013; Frakes et al. 2020; Vergani-Junior et al. 2026).

Endoplasmic reticulum quality control (ERQC) mechanisms are linked to polyQ aggregation in the cytoplasm (Duennwald and Lindquist 2008; Truttmann et al. 2018; Van Pelt and Truttmann 2025; Wei et al. 2026) as well as the inter-cellular spread of α-synuclein and tau aggregates (Wang et al. 2024). ERQC mechanisms are a critical component of the PN given that approximately one third of the proteome is synthesized at the ER (Braakman and Bulleid 2011; Martínez et al. 2017; Guay et al. 2023). The proper folding and modification of newly synthesized proteins, as well as the elimination of misfolded proteins, are mediated by various ERQC mechanisms including chaperones, ER-Associated Degradation (ERAD), the Ubiquitin Proteasome System, and autophagy (Vembar and Brodsky 2008; Araki and Nagata 2011; Martínez et al. 2017). Another critical ERQC mechanism is the ER Unfolded Protein Response (UPR^ER^), which acts as the first line of defense when unfolded or misfolded proteins accumulate in the ER (Vembar and Brodsky 2008; Powers and Balch 2013; Wang and Kaufman 2016; Hetz et al. 2020). The UPR^ER^ is comprised of three parallel signaling networks, each initiated by an ER-localized stress sensor (Walter and Ron 2011; Taylor 2016; Martínez et al. 2017; Hetz et al. 2020). Inositol-requiring enzyme-1 (IRE-1) is a stress sensor that is part of the most evolutionarily conserved branch of the UPR^ER^. IRE-1 initiates downstream signaling by splicing the *XBP1* transcription factor to generate activated XBP-1s (Walter and Ron 2011). Interestingly, several studies have linked activation of stress response pathways, including the UPR^ER^, to improvements in organismal health and longevity (Morley et al. 2002; Morley and Morimoto 2004; Steffen et al. 2008; Ben-Zvi et al. 2009; Chen et al. 2009; Henis-Korenblit et al. 2010; Taylor and Dillin 2013; Frakes et al. 2020). For example, activation of the IRE-1/XBP-1s branch of the UPR^ER^ is required for insulin/IGF1-like signaling and dietary restriction to extend lifespan in *C. elegans* (Henis-Korenblit et al. 2010; Matai et al. 2019). Additionally, expression of constitutively-active XBP-1s in *C. elegans* neurons or glial cells acts non-cell autonomously to enhance ER stress resistance in distal tissues to extend lifespan (Taylor and Dillin 2013; Frakes et al. 2020).

<u>U</u>biquitin-fold <u>m</u>odifier <u>1</u> (UFM1) is a highly conserved ubiquitin-like protein (UBL) that has emerged as a major regulator of ER protein quality control and stress responses (Hertel et al. 2013; Walczak et al. 2019; Bruck-Haimson et al. 2026; Kaiser et al. 2026; Komatsu et al. 2026; Kuang et al. 2026; Mihailovic et al. 2026). UFMylation is a post-translational modification where UFM1 is covalently attached to lysine residues on substrates using a dedicated enzyme cascade comprised of a single E1, E2, and E3 enzyme (Komatsu et al. 2004; Komatsu et al. 2024). In mammals, UFM1 is synthesized as a precursor and processed into the mature form by the proteases UFSP1 or UFSP2 to expose a C-terminal glycine residue that is covalently attached to substrates (Kang et al. 2007). Mature UFM1 undergoes trans-activation by the E1 activating enzyme UBA5 homodimer and is transferred to the catalytic cysteine on the E2 conjugating enzyme UFC1. Transfer of UFM1 from UFC1 onto target lysines on substrates requires a heterotrimer consisting of a non-canonical scaffold-type E3 ligase UFL1, an ER-anchored adaptor protein UFBP1 (Peter et al. 2022), and, in some cases, a substrate-specific adaptor CDK5RAP3 (Komatsu et al. 2026) (Supplementary Fig. 1a). Similar to other UBLs, UFM1 can be proteolytically cleaved from target substrates by the de-UFMylating proteases UFSP1 or UFSP2 (Kang et al. 2007; Komatsu et al. 2026). Although many proteins have been shown to be UFMylated (Wei and Xu 2016; Banerjee et al. 2020; Komatsu et al. 2024), it is not as clear whether these proteins are UFMylated at the ER where the UFL-1 E3 ligase complex resides, or if UFMylation of the substrate is physiologically relevant *in vivo*. One of the major proteins UFMylated at the ER is the large ribosomal subunit RPL-26 (Walczak et al. 2019; Scavone et al. 2023) where it is involved in ribosome-associated protein quality control. Loss of UFM1 pathway components in mice results in defects in several physiological processes, likely due to dysfunction in cells and tissues most sensitive to disruption of optimal ER function. For example, deletion of UFM1 pathway components in mice results in early lethality due to defects in hematopoiesis and severe anemia (Tatsumi et al. 2010; Zhang et al. 2012; Cai et al. 2015; Weisz-Hubshman et al. 2022). In *C. elegans*, loss of function of *uba-5*/UBA5 and *ufm-1*/UFM1 is not lethal but results in alterations in proteostasis and increased resistance to stress (Hertel et al. 2013; Bruck-Haimson et al. 2026; Kaiser et al. 2026). In humans, mutations and variants in UFM1 pathway components are associated with neurodevelopmental disorders such as fatal congenital neuropathy (Cabrera-Serrano et al. 2020), infantile encephalopathy (Colin et al. 2016; Muona et al. 2016; Nahorski et al. 2018) and ataxia (Duan et al. 2016). However, the physiological functions of the UFM1 pathway during stress and aging are not well understood and have only recently begun to be explored.

Here we show that loss of the UFM-1 pathway in *C. elegans* promotes activation of the UPR^ER^ under stress conditions and increases the number of protein aggregates based on a muscle-expressed YFP-tagged polyQ protein with 40 glutamines. We find that the UFM-1 pathway can act cell autonomously and non-cell autonomously to regulate muscle polyQ aggregates. The increase in aggregates observed in UFM-1 pathway mutants is mediated by the ribosomal subunit RPL-26, which is involved in ribosome-associated quality control at the ER. Physiologically, loss of the UFM-1 pathway enhances intestinal healthspan, increases survival under ER stress and suppresses the detrimental effects of polyQ expression on lifespan. Our findings indicate that the UFM-1 pathway is an important regulator of proteostasis and that inhibition of UFMylation may be beneficial for organismal health and longevity in the face of chronic proteotoxic stress.

## Materials and Methods

### Strain maintenance and genetic crosses

Worms were maintained at 20°C on standard Nematode Growth Medium (NGM) agar plates spotted with OP50 *E. coli* unless otherwise noted. 3-4 larval stage 4 (L4) worms of healthy, normal-growing strains were transferred twice a week to new plates spotted with OP50. Genotyping during genetic crosses was performed using PCR and/or sequencing to detect deletion alleles or point mutations, fluorescence to follow reporters or co-injection markers, or a combination of these approaches. A list of strains used in this study is available in Supplementary Table 1. A list of primer sequences used for PCR genotyping and sequencing is available in Supplementary Table 2.

### Molecular Biology

Plasmids were constructed with standard restriction enzyme cloning techniques. See Supplementary Table 2 for all primer sequences used for PCR and subcloning.

FJ#154 -*snb-1p::ufl-1* was constructed by using PCR to generate the *ufl-1* ORF from cDNA using forward and reverse primers engineered with flanking Nhe I and Kpn I restriction sites for subcloning. The amplified PCR product and plasmid FJ#19 (pPD49.26 backbone containing the *snb-1p* promoter) were digested with Nhe I and Kpn I and *ufl-1* cDNA was inserted. FJ#154 was injected into N2 animals at 50ng/µL with *unc-122p::GFP* (Addgene #8937) at 25ng/µL and GeneRuler 1kb Plus DNA Ladder (Thermo Scientific #SM1331) at 25ng/µL to make *pzEx505*.

FJ#155 – *myo-3p::ufl-1* was constructed by removing *pvf-1::gfp* from FJ#150 (*myo-3p::pvf-1::gfp*) (Luth et al. 2021) and inserting *ufl-1* from FJ#154 using Nhe I and Kpn I sites. FJ#155 was injected into N2 animals at 50ng/µL with *unc-122p::GFP* (Addgene #8937) at 25ng/µL and GeneRuler 1kb Plus DNA Ladder (Thermo Scientific #SM1331) at 25ng/µL to make *pzEx496*.

FJ#156 – *vha-6p::ufl-1* was constructed by removing *apm-1::mcherry* from FJ#125 (*vha-6p::apm-1:mcherry*) (Garafalo et al. 2015) and inserting *ufl-1* from FJ#154 using Nhe I and Kpn I sites. FJ#156 was injected into N2 animals at 50ng/µL with *unc-122p::GFP* (Addgene #8937) at 25ng/µL and GeneRuler 1kb Plus DNA Ladder (Thermo Scientific #SM1331) at 25ng/µL to make *pzEx504*.

### RNA Isolation and quantitative real-time PCR

Total RNA was isolated from 6-12 6cm plates per genotype of age-matched L4 animals treated with vehicle (H_2_O) or 1mM DTT in M9 buffer for 4h at 20°C by lysing in Trizol LS (Invitrogen #10296028) and extracting with the Qiagen RNeasy Fibrous Tissue Mini Kit (Qiagen #74704) with on-column DNase treatment. cDNA from the RNA preparations was synthesized using the BioRad iScript RT Reaction Kit (BioRad #1708890). RT-qPCR was performed using an Applied Biosystems StepOnePlus Real-Time PCR System (Applied Biosystems, Thermo Fisher Scientific) with Applied Biosystems PowerUp SYBR Green Master Mix (Applied Biosystems, Thermo Fisher Scientific #25741). The ΔΔCt method (Pfaffl 2001) was used to compute the relative amount of *xbp-1s* mRNA or total *xbp-1* mRNA compared to a reference gene (*act-1*). Primers used for each gene are listed in Supplementary Table 2.

### Healthspan and Lifespan assays

#### Pharyngeal Pumping assay

Pharyngeal pumping was performed as previously described (Huang et al. 2004; Laranjeiro et al. 2019). The assay was performed on L4, day 8 adult, or day 11 adult animals. To age animals, L4 animals were transferred to new plates and allowed to age to adult day 8 or adult day 11, with the day following the transfer considered as adult day 1. Animals were moved to new plates every 2-3 days to avoid starvation. Briefly, on the day of the assay, animals were transferred to new NGM plates seeded with OP50 *E. coli* and allowed to acclimate for at least 30 minutes. Following the acclimation period, 1-minute videos were obtained using an iPhone 15 Pro mounted to the dissecting microscope eyepiece using an adapter. Videos were taken at 4X magnification on the microscope and 2.4X (for day 8 and day 11 adult animals) or 3.2X (for L4 animals) on the phone. Playback speed of videos was decreased to 0.25X or 0.5X and pharyngeal pumping was counted manually for 30 seconds from each video. All video files were blinded before analysis. The results are expressed as total number of pharyngeal pumps per 30 seconds averaged across each worm of a given genotype.

#### Intestinal Barrier Integrity Assay

Intestinal barrier integrity was assayed as previously described with small modifications (McGhee 2007; Gelino et al. 2016; Laranjeiro et al. 2019). The assay was performed on L4, day 8 adult, or day 11 adult animals. To age animals, L4 animals were transferred to new plates and allowed to age to adult day 8 or adult day 11, with the day following the transfer considered as adult day 1. Animals were moved to new plates every 2-3 days to avoid starvation. Briefly, on the day of the assay, animals were transferred to a 15µL droplet containing blue dye (5% w/v Brilliant Blue FCF dye in OP50 *E. coli*) (Selleck Chemical LLC #E7158) on a coverslip. The coverslip with the droplet was placed in a 3cm plate with the lid on and placed in a box with a damp paper towel at 20°C to prevent evaporation. Animals were incubated for 3 hours. Following incubation, animals were removed from the dye and placed on new NGM plates seeded with OP50 *E. coli*. Animals were paralyzed with 2.44µM Levamisole (Tetramisole-HCl, Sigma #L9756) in a 2% agarose pad. Number of animals with and without intestinal leakage phenotypes were manually scored using a Zeiss Axioscope Imager M1 with 100X Plan-Apochromat (Numerical Aperture 1.4 Oil DIC) objective. The results are expressed as percentage of animals with intestinal leakage for each genotype.

#### DTT Survival assay

Fresh NGM plates supplemented with 10mM DTT (1M DTT stock in H_2_O, Sigma #D0632) were prepared the day before the assay. Plates were seeded with fresh OP50 *E. coli* and allowed to dry overnight. On the day of the assay, 15-20 young adult animals were transferred onto plates with DTT and incubated at 20°C. 3 plates were prepared per genotype per treatment time. Animals were scored for survival by prodding on the head with a platinum wire after 16h, 20h, and 24h of treatment. Animals were considered dead if they failed to respond to prodding. 3-7 independent trials were carried out for each genotype. Results are expressed as percent survival for each genotype at each time point.

#### Lifespan assay

Lifespan assays were performed as previously described, with some modifications (Sutphin and Kaeberlein 2009; Amrit et al. 2014). Briefly, 5-10 gravid adults were allowed to egg lay for 2-4 hours on seeded NGM plates and then removed. Hatched animals were scored for survival and transferred to fresh seeded NGM plates every 1-2 days. 80-120 animals were used for each genotype per experiment and 3-6 independent experiments were carried out. Animals were considered dead if they failed to respond to prodding. Animals were eliminated from the experiment if they crawled off the plate or internal egg hatching (bagging) occurred.

### Epifluorescent imaging

All *unc-54p::Q40::YFP* (*rmIs133*) animals were imaged using a Zeiss Axioscope Imager M1 with a GFP filter cube and equipped with an Orca-ER charge-coupled device camera (Hamamatsu) and MetaMorph, version 7.1 software (Molecular Devices). Worms were immobilized on a 2% agarose pad containing 2.4mM levamisole (Tetramisole-HCl, Sigma #L9756) in a 5µL drop of M9 Buffer. A glass coverslip was inverted over the agarose pad. All imaging was performed on L4.4 stage worms to minimize variation in aggregate number between animals at different substages (Mok et al. 2015). Vulval staging was determined using the 63X Plan-Apochromat objective (Numerical Aperture 1.4 Oil DIC). Fluorescence images were acquired at 5X magnification (5X A-Plan objective, Numerical Aperture 0.12 Air) using the GFP filter cube with the following settings: Exposure time: 200ms; Transmission: 25%; Binning: 1; Gain: 0; Z-Stack: 11 slices (40µm). All images were saved as TIFF files.

### Confocal imaging

Images were captured using a Zeiss LSM800 confocal microscope and Zen Blue software (Zeiss). Imaging was performed on L4 stage animals. Worms were immobilized on a glass coverslip for 5 minutes in a 5 µL drop of M9 Buffer containing 30mg/mL BDM (2,3-Butanedione monoxime, Sigma #B0753) unless otherwise stated. A 2% agarose pad on a glass slide was inverted over the droplet to mount the worms on the slide. All images were saved as CZI files. Specific Acquisition settings are described in detail below.

#### Heat Shock

For heat-shock experiments, L4 animals were transferred to fresh NGM plates and heat shocked at 30°C for 4 hours. Plates were staggered to allow for sufficient imaging time between genotypes. Only L4 animals were imaged. Slides were prepared as described above.

#### unc-54p::Q24::YFP imaging

Fluorescence images were acquired at 20X magnification (20X Plan-Apochromat objective, Numerical Aperture 0.8 Air) using the following settings: Acquisition Area: 638.6µm X 638.6µm; Pinhole: 30µm; EGFP Channel: 488nm laser: 0.8%, Gain: 500V; Z-Stack: 21 slices (40µm).

#### xbp1p::xbp-1s::GFP imaging

Fluorescence and T-PMT images were acquired at 63X magnification (63X Plan-Apochromat objective, Numerical Aperture 1.4 Oil DIC) from the mid-intestine centered on the vulva for each animal using the following settings: Acquisition Area: 101.41µm X 101.41µm; Pinhole: 41µm; EGFP Channel: 488nm laser: 2.0%, Gain: 500V; T-PMT Channel: Gain 300V; Z-Stack: 32 slices (20.15µm).

#### nhx-2p::CPL-1*::YFP (nhx-2p::CPL-1^W32A, Y35A^::YFP) imaging

Fluorescence and T-PMT images were acquired at 20X magnification (20X Plan-Apochromat objective, Numerical Aperture 0.8 Air) using the following settings: Acquisition Area: 638.9µm X 638.9µm; Pinhole: 30µm; EGFP Channel: 488nm laser: 3.5% (*sel-11* mutants were imaged at 1%), Gain: 500V; T-PMT Channel: Gain 225V; Z-Stack: 25 slices (30µm).

#### sqst-1p::sqst-1::GFP imaging

Fluorescence and T-PMT images were acquired at 63X magnification (63X Plan-Apochromat objective, Numerical Aperture 1.4 Oil DIC) from the anterior intestine behind the pharynx and the mid-intestine centered on the vulva for each animal using the following settings: Acquisition Area: 101.41µm X 101.41µm; Pinhole: 45µm; EGFP Channel: 488nm laser: 9.0%, Gain: 700V; T-PMT Channel: Gain 250V; Z-Stack: 32 slices (20.15µm).

#### Fluorescence Recovery After Photobleaching (FRAP)

Fluorescence images were acquired at 63X magnification (63X Plan-Apochromat objective, Numerical Aperture 1.4 Oil DIC). Worms were immobilized using a combination of levamisole and polystyrene microspheres. Briefly, animals were placed in a 5µL drop of 0.1µm Polybead polystyrene microspheres (Polysciences, Inc. #00876) on a glass coverslip and covered with a 3% agarose pad containing 2.4mM freshly prepared levamisole (Tetramisole-HCl, Sigma #L9756). One aggregate was randomly selected for analysis for each animal imaged. Images were acquired using the following settings: Acquisition Area: 101.41µm X 101.41µm; Pinhole: 45µm; EGFP Channel: 488nm laser: 0.2%, Gain: 530V. The midpoint of each aggregate was determined and used as the center of all z-stack acquisitions within the time-series. The Z-stacks consisted of 32 slices spanning 20.15µm. Prior to photobleaching, 6 pre-bleach images were acquired from the midpoint of the aggregate over 12.29s. Half of the aggregate was photobleached using 2 iterations of the 488nm laser at 50% laser power to achieve ∼80% bleaching. Immediately following photobleaching, a z-stack spanning the entire aggregate was acquired (“0-min post-bleach”). Additional z-stacks were acquired at 2, 4, 6, 8, and 10-minutes post-bleach using identical imaging parameters. Only aggregates in which bleaching extended through the whole aggregate volume were included in the analysis.

### Image analysis

Maximum intensity projections of z-series stacks were made using FIJI and background subtracted using the rolling ball algorithm (radius = 50 pixels). If multiple channels were acquired, images were split into individual channels using FIJI before making maximum intensity projections and background subtraction. Specific analysis methods are described in detail below.

#### unc-54p::Q40::YFP image analysis

All images were analyzed using custom FIJI scripts, followed by a custom MATLAB script. A region of interest (ROI) encompassing the entire animal was defined manually. The Otsu thresholding method was used to define aggregate identity and minimum and maximum thresholds were empirically determined for each day of imaging using representative images. Thresholding parameters were subsequently uniformly applied to all images for each genotype imaged on that day.

Aggregate size and number were quantified using the Analyze Particles function in FIJI and restricted to the manually defined ROI. Output files were generated for each image containing the number of aggregates counted and area measurements for each animal. The output files were then run through a custom MATLAB script to extract aggregate number per animal, which are the values used for downstream graphical presentation and statistical analyses.

#### unc-54p::Q24::YFP image analysis

A ROI encompassing the entire animal was defined manually. Area and raw integrated fluorescence density were measured using FIJI. Fluorescence intensity was normalized to animal area. Each value was normalized to the wild-type average for that imaging session. These normalized values were used for downstream graphical presentation and statistical analyses.

#### xbp-1p::xbp-1s::GFP image analysis

ROIs were manually drawn around all visible intestinal nuclei within each animal. Area and raw integrated fluorescence density were measured using FIJI. Fluorescence intensity was normalized to intestinal nuclear area and then averaged for all nuclei within an individual animal to generate a single value per animal. Each value was normalized to the wild-type average for that imaging session. These normalized values were used for downstream graphical presentation and statistical analyses.

#### nhx-2p::CPL-1*::YFP (nhx-2p::CPL-1^W32A, Y35A^::YFP) image analysis

A segmented line ROI (line thickness = 172 pixels) was manually drawn on the brightfield image beginning immediately behind the pharynx and extending the length of the animal to the tail. The ROI was transferred to the corresponding GFP image, and area and raw integrated fluorescence density were measured using FIJI. Fluorescence intensity was normalized to animal area. Each value was normalized to the wild-type average for that imaging session. These normalized values were used for downstream graphical presentation and statistical analyses.

#### sqst-1p::sqst-1::GFP image analysis

A freehand ROI was manually drawn on the brightfield image encompassing the entire portion of the intestine in the image. The ROI was transferred to the corresponding GFP image, and area and raw integrated fluorescence density were measured using FIJI. Fluorescence intensity was normalized to animal area. Each value was normalized to the wild-type average for that imaging session. These normalized values were used for downstream graphical presentation and statistical analyses.

#### Fluorescence Recovery After Photobleaching (FRAP)

FRAP image analysis was performed using FIJI. Maximum intensity projections were generated for each post-bleach time point z-series stack. Images were screened prior to analysis to confirm that photobleaching extended throughout the entire aggregate volume. Aggregates that were not completely bleached were excluded from analysis. A new image stack containing the 6 pre-bleach images, the bleach image, and the 6 post-bleach maximum intensity projections was generated for each aggregate. The TurboReg plugin (FIJI) was used to correct for image drift.

Three ROIs were manually drawn for each image: one encompassing the bleached portion of the aggregate, one encompassing the unbleached reference region of the same aggregate, and a nearby background region. Integrated density measurements were collected for each ROI at every time point. Background fluorescence was subtracted from both the bleached and reference ROIs to obtain background corrected fluorescence values. The background corrected fluorescence values from the bleached ROI were corrected for potential photobleaching during the acquisition by using the background corrected unbleached reference ROI fluorescence values and then normalized to the average pre-bleach fluorescence intensity for each aggregate.

Normalized fluorescence recovery values were scaled such that the mean pre-bleach fluorescence intensity was set to 1 and the fluorescence intensity immediately following photobleaching was set to 0. Fluorescence recovery curves were made by graphing normalized fluorescence recovery as a function of time after photobleaching.

### Statistical Analysis

Each data point represents a single animal in all imaging experiments unless otherwise stated. For normalized datasets, measurements obtained from animals imaged across different experimental days were normalized to the average value of the corresponding wild-type animals imaged on the same day prior to pooling across biological replicates. All data are presented as mean ± SEM unless otherwise stated.

All datasets were assessed for normality using the D’Agostino-Pearson normality test or the Shapiro-Wilk normality test, followed by analysis using the appropriate parametric or non-parametric statistical tests. All statistical analyses were performed in GraphPad Prism version 11. The specific statistical tests used, sample sizes, and significance levels are reported in the corresponding figure legends.

## Results

### Loss of the UFM-1 pathway promotes activation of the UPR^ER^ under stress conditions

Prior studies showed that loss of the UFM1 pathway leads to activation of the UPR^ER^ in mammalian cells and *C. elegans* (Zhang et al. 2012; Hertel et al. 2013; Cai et al. 2015; Kaiser et al. 2026). Disruption of the UFM1 pathway in mammalian cells causes increased expression of several proteins involved in UPR^ER^ activation and signaling, including Grp78/Bip (Zhang et al. 2012; Cai et al. 2015). In *C. elegans*, loss-of-function mutations in *ufm-1* or the UFM-1 pathway E1 enzyme *uba-5*, results in low UPR^ER^ activation in the intestine under basal conditions and enhanced UPR^ER^ activation under conditions that cause proteotoxic stress (i.e., DTT and heat) based on a *hsp-4p::gfp* transcriptional reporter (Bip reporter) (Hertel et al. 2013; Kaiser et al. 2026). However, a recent study in *C. elegans* showed that RNAi-knockdown of *ufm-1* has no effect on UPR^ER^ activation based on the same *hsp-4p::gfp* reporter in *C. elegans* (Bruck-Haimson et al. 2026).

Given these conflicting results, we tested if loss of the UFM-1 pathway could activate the UPR^ER^ using a distinct fluorescent reporter that directly measures *xbp-1* splicing. This reporter expresses genomic *xbp-1* and GFP under control of the *xbp-1* promoter such that GFP is only in-frame with the XBP-1s coding region when the UPR^ER^ is activated and IRE-1 removes an intron upstream of GFP (Shim et al. 2004; Özbey et al. 2020). Thus, XBP-1s::GFP intensity in nuclei of the intestine can be used to measure *xbp-1s* splicing as a readout for UPR^ER^ activation. We first tested whether loss-of-function mutations in the UFM-1 pathway E1 enzyme *uba-5 (uba-5(ok3364)*) or E3 ligase *ufl-1* (*ufl-1(tm5599)*) activated the UPR^ER^ in larval stage 4 (L4) animals under basal conditions. The *uba-5(ok3364)* allele is a functional null as it is missing the active site cysteine (Cys260) and is unable to bind UFM-1 (Hertel et al. 2013) (Supplementary Fig. 1b). The *ufl-1(tm5599)* mutant allele is comprised of a 696 bp deletion from the 4^th^-6^th^ exon, likely resulting in a non-functional ligase (Supplementary Fig. 1b). Under basal conditions, we found no change in XBP-1s::GFP fluorescence intensity in nuclei in the intestine of *uba-5(ok3364)* or *ufl-1(tm5599)* mutants compared to wildtype (Fig. 1a and b, 1^st^ vs 2^nd^ and 3^rd^ bars). Interestingly, although XBP-1s::GFP intensity did not increase in wild-type animals exposed to a low level of heat-shock stress (Fig. 1a and 1b, 1^st^ vs 4^th^ bars), XBP-1s::GFP intensity did increase in both *uba-5(ok33654)* and *ufl-1(tm5599)* mutants following heat-shock stress compared to wildtype at baseline (Fig. 1a and 1b, 1^st^ vs 5^th^ and 6^th^ bars). These results suggest that loss of the UFM-1 pathway promotes activation of the UPR^ER^ after exposure to heat-shock stress.

**Fig. 1.**
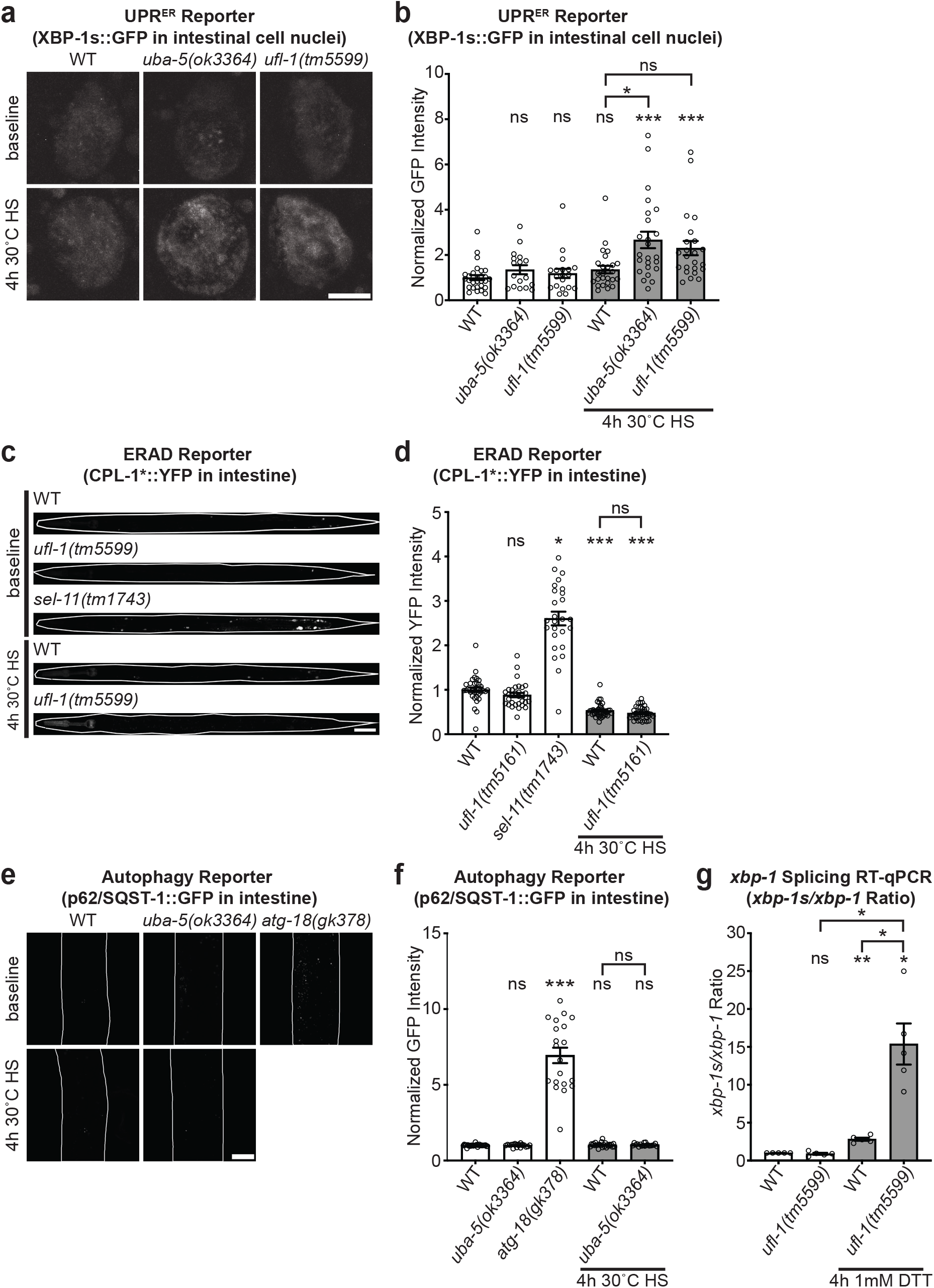
Loss of the UFM-1 pathway promotes activation of the UPR^ER^ under proteotoxic stress. (a) Representative images of XBP-1s::GFP (*rmsIs8*), an UPR^ER^ splicing reporter, in intestinal cell nuclei at baseline and following heat shock (HS, 4h at 30°C) for wild-type, *uba-5(ok3364)*, and *ufl-1(tm5599)* animals. Scale bar represents 5µm. (b) Quantification of GFP fluorescence intensity of XBP-1s::GFP in intestinal cell nuclei at baseline and after heat shock (HS, 4h at 30°C) for wild-type (baseline, n=28; HS, n=26), *uba-5(ok3364)* (baseline, n=17; HS, n=24), and *ufl-1(tm5599)* (baseline, n=19; HS, n=23) animals. Kruskal-Wallis test with Dunn’s Multiple Comparisons test revealed a significant difference between: *ufl-1(tm5599)* (p = 0.0002) or *uba-5(ok3364)* (p < 0.0001) after heat shock compared to wildtype at baseline, and *uba-5(ok3364)* (p = 0.0321) after heat-shock compared to wildtype after heat shock. (c) Representative images of CPL-1*::YFP (*hhIs113*), an ERAD reporter, at baseline and after heat-shock (HS, 4h at 30°C) for wildtype and *ufl-1(tm5161)*, and *sel-11(tm1743)* at baseline. *sel-11(tm1743)* was included as a positive control. *sel-11(tm1743)* images were taken at lower laser intensity than other samples (see Materials and Methods). All animals were straightened and anterior is oriented to the left of the image. Some bleed-through of the co-injection marker *myo2p::mCherry* (expressed in the pharynx) is visible in the GFP channel in some images. Scale bar represents 50µm. (d) Quantification of fluorescence intensity of CPL-1*:YFP at baseline and after heat shock (HS, 4h at 30°C) for wildtype (baseline, n=34; HS, n=37) and *ufl-1(tm5161)* (baseline, n=32; HS, n=39). *sel-11(tm1743)* (baseline, n=26) is a positive control. Kruskal-Wallis test with Dunn’s Multiple Comparisons revealed a significant difference between wildtype (p < 0.0001) and *ufl-1(tm5161)* (p < 0.0001) after heat shock compared to wildtype at baseline, and between *sel-11(tm1743)* (p = 0.0183) and wildtype at baseline. (e) Representative images of SQST-1::GFP (*bpIs151*), an autophagy reporter, at baseline and after heat-shock (HS, 4h at 30°C) for wildtype and *uba-5(ok3364)*, and *atg-18(gk378)* at baseline as a positive control. The animal is outlined in white. Scale bar represents 20µm. (f) Quantification of GFP fluorescence intensity of SQST-1::GFP at baseline and after heat-shock (HS, 4h at 30°C) for wildtype (baseline, n=29; HS, n=26), *uba-5(ok3364)* (baseline, n=20; HS, n=19), and *atg-18(gk378)* (baseline, n=20) animals. *atg-18(gk378)* was included as a positive control. Kruskal-Wallis test with Dunn’s Multiple Comparisons test revealed no significant differences between wild-type (p > 0.9999) and *ufl-1(tm5599)* (p > 0.9999) animals after heat-shock compared to wildtype at baseline but did reveal a significance difference between *atg-18(gk378)* (p < 0.0001) and wildtype at baseline. (g) Ratio of relative expression of *xbp-1s* mRNA to total *xbp-1* mRNA (normalized to *act-1*) under baseline and DTT-treated conditions (1mM DTT for 4h) measured by RT-qPCR for wild-type and *ufl-1(tm5599)* animals. n = 5 independent trials, each data point represents the average of 2-3 technical replicates within a trial. Brown-Forsythe and Welch ANOVA tests with Dunnett’s Multiple Comparisons test revealed a significant difference between wildtype (p = 0.0038) and *ufl-1(tm5599)* (p = 0.0258) after DTT treatment compared to wildtype at baseline, *ufl-1(tm5599)* (p = 0.0252) animals after DTT treatment compared to *ufl-1(tm5599)* at baseline, and *ufl-1(tm5599)* (p = 0.0415) after DTT treatment compared to wildtype after DTT treatment. Unless otherwise noted, all graphs show mean ± SEM. Differences versus wild-type baseline are shown above each bar. All other comparisons are marked by brackets. ns, p > 0.05; *, p < 0.05; **, p < 0.01; ***, p < 0.001.

We next tested whether loss of the UFM-1 pathway affects two other proteostatic pathways, ER Associated Degradation (ERAD) and autophagy. We measured ERAD function by monitoring the accumulation of a model ERAD client: a YFP-tagged, misfolding-prone version of lysosomal cathepsin L-like peptidase (CPL-1*::YFP) expressed specifically in the intestine (under control of *nhx-2p*). Under basal conditions, misfolded CPL-1*::YFP is degraded via ERAD. However, disruption of ERAD results in increased fluorescence of CPL-1*::YFP due to the accumulation of misfolded CPL-1*::YFP in the ER (Miedel et al. 2012; Finger et al. 2019) (Fig. 1c). As expected, loss of the ERAD E3 ligase *sel-11*/HRD-1 disrupts ERAD and results in the accumulation of CPL-1*::YFP in the ER (Fig. 1c and d, 1^st^ vs 3^rd^ bars). We found that *ufl-1(tm5161)* mutants do not affect CPL-1*::YFP fluorescence under basal conditions (Fig. 1c and d, 1^st^ vs 2^nd^ bar), suggesting that ERAD is not blocked or triggered in the absence of *ufl-1*. We next tested if the same heat-shock treatment described above would affect ERAD function. We observed that a low level of heat-shock results in decreased CPL-1*::YFP accumulation in wild-type animals compared to baseline (Fig. 1c and d, 1^st^ vs 4th bars). This result might be consistent with heat-shock causing an increase in CPL-1*::YFP misfolding followed by increased degradation via ERAD. We found that CPL-1*::YFP fluorescence decreased to the same extent in *ufl-1(tm5161)* mutants under these heat-shock conditions (Fig. 1c and d, 4^th^ vs 5^th^ bar), suggesting that loss of the UFM-1 pathway does not have a major effect on ERAD either with or without heat-shock stress.

We measured autophagy activation by monitoring the accumulation of a fluorescently-tagged autophagy receptor, p62/SQST-1::GFP, under control of its own promoter (*sqst-1p*). Under basal conditions, SQST-1::GFP is diffusely localized in the cytoplasm and expressed at a low level due to continuous degradation via autophagy. However, when autophagy is impaired, p62/SQST-1::GFP protein accumulates in puncta in various tissues including the intestine (Tian et al. 2010; Guo et al. 2014). As expected, we found that p62/SQST-1::GFP fluorescence increases in the intestine in animals lacking the central autophagy regulator *atg-18* (Fig. 1e and f, 1^st^ vs 3^rd^ bars). However, we observed no change in p62/SQST-1::GFP accumulation in *uba-5(ok3364)* loss-of-function mutants under basal conditions or after heat-shock stress (Fig. 1e and f, 1^st^ vs 2^nd^ bars, 4^th^ vs 5^th^ bars). Taken together, our results show that loss of the UFM-1 pathway does not have a major effect on the UPR^ER^ under basal conditions but does promote activation of the UPR^ER^ after heat-shock stress. In addition, loss of the UFM-1 pathway does not have a major effect on activation of ERAD or autophagy with or without heat-shock stress based on the CPL-1*::YFP and p62/SQST-1::GFP reporters analyzed. Our data show that among the proteostatic pathways examined, the UPR^ER^ exhibited the most robust activation in response to loss of the UFM-1 pathway.

We next used RT-qPCR to monitor activation of the UPR^ER^ in whole animals by measuring endogenous levels of spliced *xbp-1 (xbp-1s)* transcript levels in age-matched L4 animals in the absence or presence of DTT, which causes protein misfolding and proteotoxic stress. We observed no difference in the ratio of *xbp-1s* to total *xbp-1* (*xbp-1s*/*xbp-1*) transcripts in *ufl-1(tm5599)* mutants compared to wild-type controls at baseline (Fig. 1g, 1^st^ vs 2^nd^ bar). As expected, DTT treatment resulted in an increase in the *xbp-1s*/*xbp-1* ratio (∼3-fold) in wild-type animals compared to baseline (Fig. 1g, 1^st^ vs 3^rd^ bar). Consistent with our results above using the UPR^ER^ fluorescent reporter (*XBP-1s::GFP*), we found that loss of the UFM-1 pathway in *ufl-1(tm5599)* mutants resulted in a 5-fold increase in the *xbp-1s*/*xbp-1* ratio (∼5-fold) following DTT treatment compared to wild-type animals (Fig. 1g, 3^rd^ vs 4^th^ bar). Collectively, our data show that loss of the UFM-1 pathway enhances activation of the UPR^ER^ under conditions of proteotoxic stress.

### Loss of the UFM-1 pathway results in increased polyQ aggregates in muscle

To further investigate the effects of loss of UFM-1 pathway genes on proteostasis, we utilized a well-characterized *C. elegans* polyglutamine (polyQ) aggregation model to monitor and assess proteostasis and cellular dysfunction (Morley et al. 2002; Nollen et al. 2004; Gidalevitz et al. 2006; Garcia et al. 2007; De-Souza et al. 2023). In this model, yellow fluorescent protein is tagged with 40 glutamines (Q40::YFP) and expressed in body wall muscle (under control of the *unc-54*/myosin promoter) (Morley et al. 2002). Q40::YFP aggregates are highly sensitive to perturbations in protein homeostasis and as a result the number of aggregates increases in response to proteotoxic stress (Morley et al. 2002; Nollen et al. 2004; Gidalevitz et al. 2006; Garcia et al. 2007; De-Souza et al. 2023) and with age, as is observed in polyglutamine aggregation diseases in humans (Morley et al. 2002). Furthermore, expression of expanded polyQ proteins in *C. elegans* imposes a chronic proteotoxic challenge to the PN, leading to reductions in healthspan and lifespan (Morley et al. 2002; Gidalevitz et al. 2006).

We used loss-of-function mutants of each component of the UFM-1 pathway to test how loss of these components would affect Q40::YFP aggregates. Because the number of Q40::YFP aggregates is sensitive to age, we only imaged L4 animals that were precisely age-matched at the L4.4 stage based on vulval morphology (see Materials and Methods). We found that loss of the UFM-1 E3 ligase *ufl-1*, using two independent loss-of-function alleles (*tm5599* and *tm5161*) (see Supplementary Fig. 1b for molecular lesions), results in an increase in the number of Q40::YFP aggregates in muscle (Fig. 2a and b, 1^st^ vs 2^nd^ and 3^rd^ bars). We then tested if loss of the other core pathway enzymes required for UFMylation were also involved in regulating polyQ aggregates. We found that loss of the E1 enzyme *uba-5* or the E2 enzyme *ufc-1* also results in an increase in the number of Q40::YFP aggregates, suggesting that loss of the core enzymes required for UFMylation alter polyQ aggregates (Fig. 2a and b, 1^st^ vs 4^th^ and 5^th^ bars). Similarly, we found that loss of the de-UFMylating protease *ufsp-2* also results in increased Q40::YFP aggregates (Fig. 2a and b, 1^st^ vs 7^th^ bar). Although counter-intuitive, this result is consistent with the additional role of UFSP-2 in processing UFM-1 precursors to the mature form that can be conjugated to target proteins (Kang et al. 2007; Hertel et al. 2013; Ha et al. 2016). Additionally, we found that loss of the pathway’s ER-anchored proteins UFBP-1, which is an essential part of the UFL-1 E3 ligase complex required for UFMylation, and ODR-4, which recruits UFSP-2 to the ER membrane, results in a similar increase in the number of Q40::YFP aggregates (Fig. 2a and b, 1^st^ vs 6^th^ or 8^th^ bars), suggesting that the UFM-1 pathway functions at the ER membrane to regulate polyQ aggregates in the cytoplasm.

**Fig. 2.**
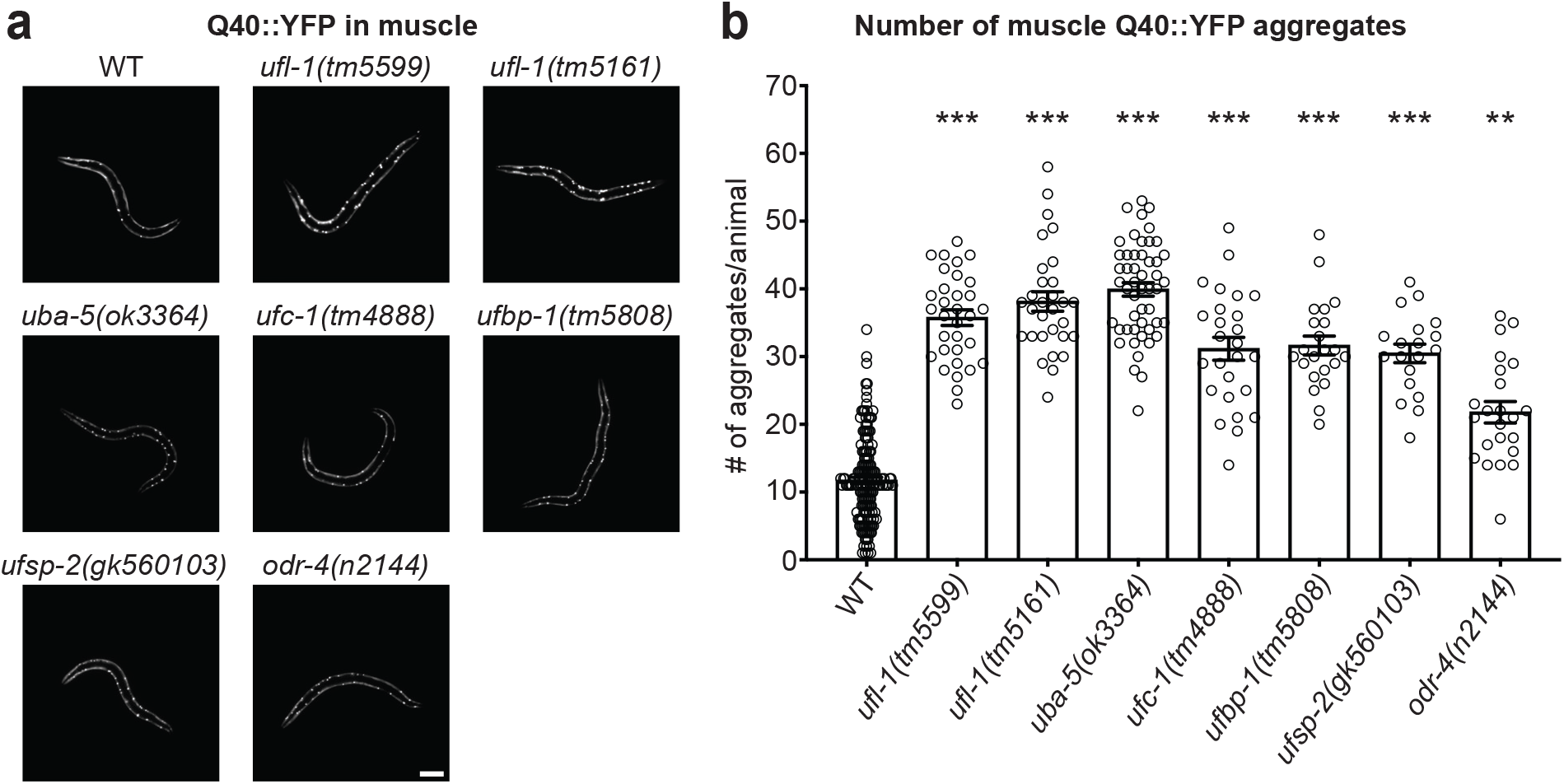
Loss of UFM-1 pathway genes leads to an increased number of polyQ aggregates in muscle. (a) Representative images of Q40::YFP in muscle (*rmIs133*) in wild-type, *ufl-1(tm5599)*, *ufl-1(tm5161)*, *uba-5(ok3364)*, *ufc-1(tm4888)*, *ufbp-1(tm5808)*, *ufsp-2(gk560103)*, and *odr-4(n2144)* animals. Scale bar represents 100µm. (b) Quantification of the number of Q40::YFP aggregates per animal in age-matched L4 wild-type (Q40::YFP in muscle) (n=169), *ufl-1(tm5599)* (n=32), *ufl-1(tm5161)* (n=30), *uba-5(ok3364)* (n=50), *ufc-1(tm4888)* (n=26), *ufbp-1(tm5808)* (n=22), *ufsp-2(gk560103)* (n=19), and *odr-4(n2144)* (n=23) animals. Kruskal-Wallis test with Dunn’s Multiple Comparisons Test revealed a significant difference between all genotypes compared to wildtype (*ufl-1(tm5599)*, p < 0.0001; *ufl-1(tm5161)*, p < 0.0001; *uba-5(ok3364)*, p < 0.0001; *ufc-1(tm4888)*, p < 0.0001; *ufbp-1(tm5808)*, p < 0.0001; *ufsp-2(gk560103)*, p < 0.0001; and *odr-4(n2144)*, p = 0.0018). Unless otherwise noted, all graphs show mean ± SEM. Differences versus wildtype are shown above each bar. **, p < 0.01; ***, p < 0.001.

We next tested whether the increase in Q40::YFP number in UFM-1 pathway mutants was due to increased expression of the transgene, which is driven by the *unc-54*/myosin promoter. To monitor *unc-54* promoter activity, we measured the expression of Q24::YFP in muscle using a transgene that is under control of the same *unc-54*/myosin promoter. Q24::YFP is below the threshold for polyQ toxicity and remains in a diffuse state in the cytoplasm even in the presence of aggregation modifiers that induce premature aggregation of transgenes with longer glutamine repeats (Garcia et al. 2007). We found that loss of *uba-5* (E1 enzyme) or *ufl-1* (E3 enzyme) had no effect on Q24::YFP fluorescence intensity compared to wild-type animals (Supplementary Fig. 2a and b), suggesting that the increase in number of Q40::YFP aggregates in UFM-1 pathway mutants cannot be attributed to increased *unc-54* promoter activity and is likely due to a disruption in protein homeostasis. Taken together, our data suggest that loss of each individual component of the UFM-1 pathway, including the core enzymes required for UFMylation and the ER membrane proteins that anchor the UFMylation machinery to the ER, leads to increased numbers of polyQ protein aggregates.

### *ufl-1* acts cell autonomously and non-cell autonomously to regulate polyQ aggregates in muscle

Because organismal proteostasis is coordinated by inter-tissue signals (Henis-Korenblit et al. 2010; Taylor and Dillin 2013; Frakes et al. 2020; Hoppe and Cohen 2020; Özbey et al. 2020) and studies have suggested that UFM-1 pathway genes are most highly expressed in the intestine (Hertel et al. 2013; Kaiser et al. 2026), we performed tissue-specific rescue experiments to test if the UFM-1 pathway regulates polyQ aggregates in muscle cell autonomously or non-cell autonomously. Surprisingly, we found that the increased number of polyQ aggregates observed in the muscle of *ufl-1(tm5599)* mutants is not rescued by restoring wild-type *ufl-1* expression in the intestine (using *vha-6p*) (Fig. 3a and b, 2^nd^ vs 3^rd^ bars). However, expression of wild-type *ufl-1* specifically in muscle (using *myo-3p*) or neurons (using *snb-1p*) was able to restore polyQ aggregate number back to baseline (Fig. 3a and b, 2^nd^ vs 4^th^ or 5^th^ bars). These data suggest that the UFM-1 pathway acts cell autonomously in muscle and non-cell autonomously from neurons to regulate polyQ aggregates in muscle and reveals the existence of an inter-tissue signal from neurons to muscle that regulates proteostasis.

**Fig. 3.**
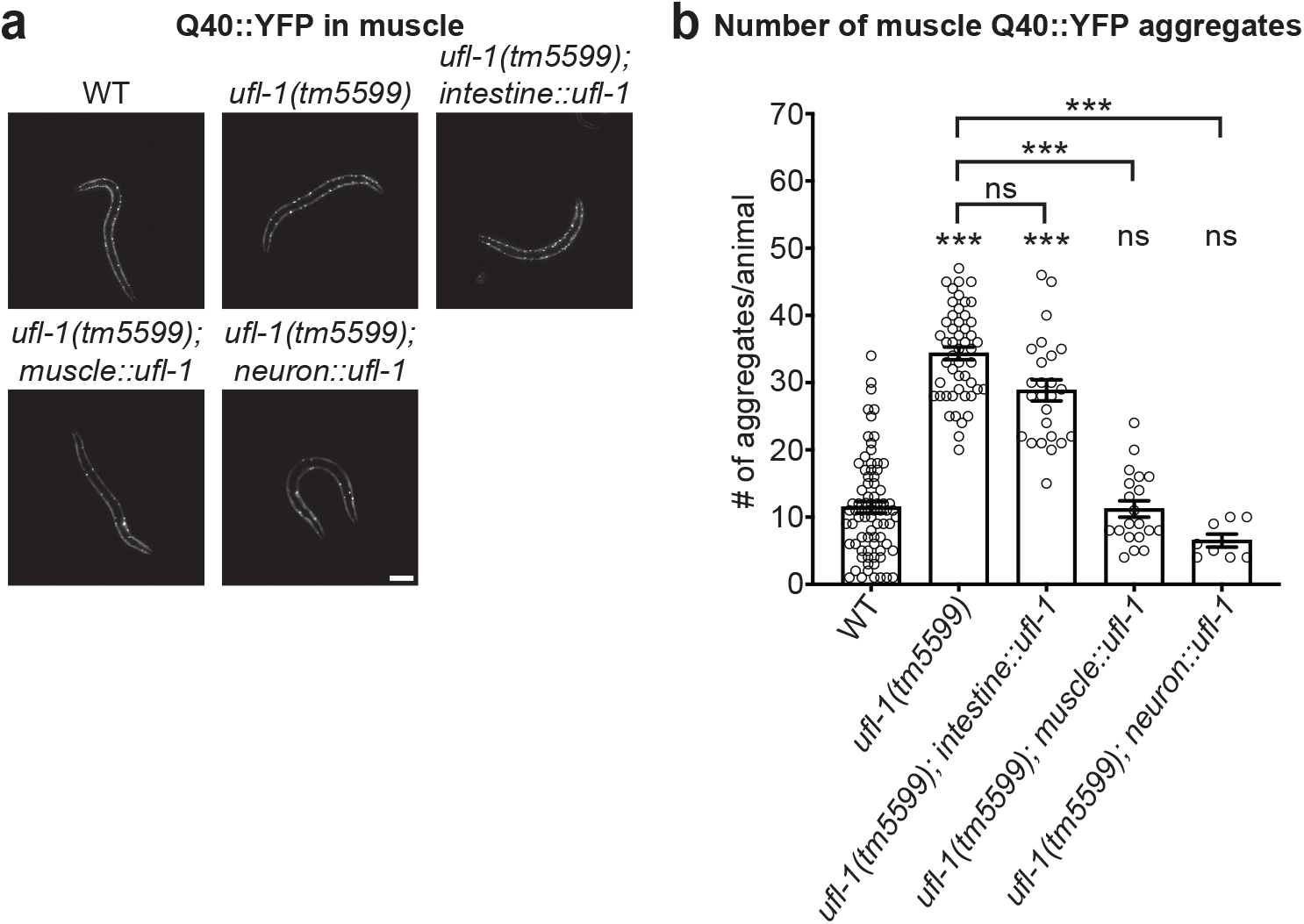
*ufl-1* acts in muscle and neurons to regulate polyQ aggregates in muscle. (a) Representative images of Q40::YFP in muscle (*rmIs133*) in wild-type, *ufl-1(tm5599); muscle::ufl-1* rescue, *ufl-1(tm5599); intestine::ufl-1* rescue, and *ufl-1(tm5599); neuron::ufl-1* rescue animals. Images were enhanced for display purposes. Scale bar represents 100µm. (b) Quantification of the number of Q40::YFP aggregates per animal in age-matched L4 wild-type (Q40::YFP in muscle) (n=77), *ufl-1(tm5599)* (n=50), *ufl-1(tm5599); muscle::ufl-1* rescue (n=20), *ufl-1(tm5599); intestine::ufl-1* rescue (n=25), and *ufl-1(tm5599); neuron::ufl-1* rescue (n=8) animals. Kruskal-Wallis test with Dunn’s Multiple Comparisons Test revealed a significant difference between: wildtype and *ufl-1(tm5599)* (p < 0.0001), wildtype and *ufl-1(tm5599); intestine::ufl-1* (p < 0.0001), *ufl-1(tm5599)* and *ufl-1(tm5599); muscle::ufl-1* (p < 0.0001), and *ufl-1(tm5599)* and *ufl-1(tm5599); neuron::ufl-1* (p < 0.0001). Unless otherwise noted, all graphs show mean ± SEM. Differences versus wildtype are shown above each bar. All other comparisons are marked by brackets. ns, p > 0.05; ***, p < 0.001.

### UFL-1-regulation of polyQ aggregates is mediated by the major UFMylation target RPL-26

Several proteins have been proposed as UFMylation substrates (Wei and Xu 2016; Walczak et al. 2019; Banerjee et al. 2020; Witting and Mulder 2021; Bruck-Haimson et al. 2026; Komatsu et al. 2026). A rigorous mass spectrometry study of endogenously-expressed proteins in mammalian cells identified RPL26 as a major UFMylation target (Walczak et al. 2019; Scavone and Kopito 2026). RPL26 is a large ribosomal subunit that is involved in ribosome quality control. UFMylation of RPL26 promotes the clearance of partially translated proteins at the ER and recycling of ribosomes (Walczak et al. 2019; Wang et al. 2020; Scavone et al. 2023; DaRosa et al. 2024; Makhlouf et al. 2024). Because our data suggest that the UFM-1 pathway acts at the ER to regulate Q40::YFP aggregates (Fig 2) and mass spectrometry data show that RPL-26 is also UFMylated in *C. elegans* (Bruck-Haimson et al. 2026), we investigated whether RPL-26 mediates the increase in Q40::YFP aggregates observed in UFM-1 pathway mutants. Mammalian RPL26 is UFMylated at K132 and K134 and mutation of these sites to arginine eliminates UFMylation (Walczak et al. 2019). Both lysines are conserved in *C. elegans* as K133 and K135 (Fig. 4a). Both of these critical lysines were mutated to arginine (K133R/K135R) in the endogenous *C. elegans rpl-26* gene using CRISPR/Cas9 gene editing to test the effect of blocking these conserved RPL-26 UFMylation sites on Q40::YFP aggregates. Interestingly, we found a significant increase in the number of Q40::YFP aggregates in *rpl-26* (K133R/K135R) mutants (*rpl-26(KK>RR))* compared to wild-type animals (Fig. 4b and c, 1^st^ vs 3^rd^ bars), and this increase was similar to the number of Q40::YFP aggregates observed in *ufl-1(tm5599)* loss-of-function mutants (Fig. 4b and c, 2^nd^ vs 3^rd^ bars). These data indicate that blocking the putative UFMylation sites on RPL-26 is sufficient to increase the number of Q40::YFP aggregates. We performed double mutant analysis to test if *rpl-26(KK>RR)* mediates the effects of *ufl-1(tm5599)* on the number of Q40::YFP aggregates. If UFMylation of these lysine residues on RPL-26 mediate the effects of UFL-1 on Q40::YFP aggregates, then we would expect *rpl-26(KK>RR); ufl-1(tm5599)* to have a non-additive effect on the number of Q40::YFP aggregates. Consistent with this idea, we found that the number of Q40::YFP aggregates in *rpl-26(KK>RR); ufl-1(tm5599)* double mutants was similar to that observed in either single mutant (Fig. 4b and c, 2^nd^ vs 4^th^ bars and 3^rd^ vs 4^th^ bars). This non-additive effect supports the idea that UFMylation of RPL-26 mediates the effects of the UFM-1 pathway on polyQ protein aggregates.

**Fig. 4.**
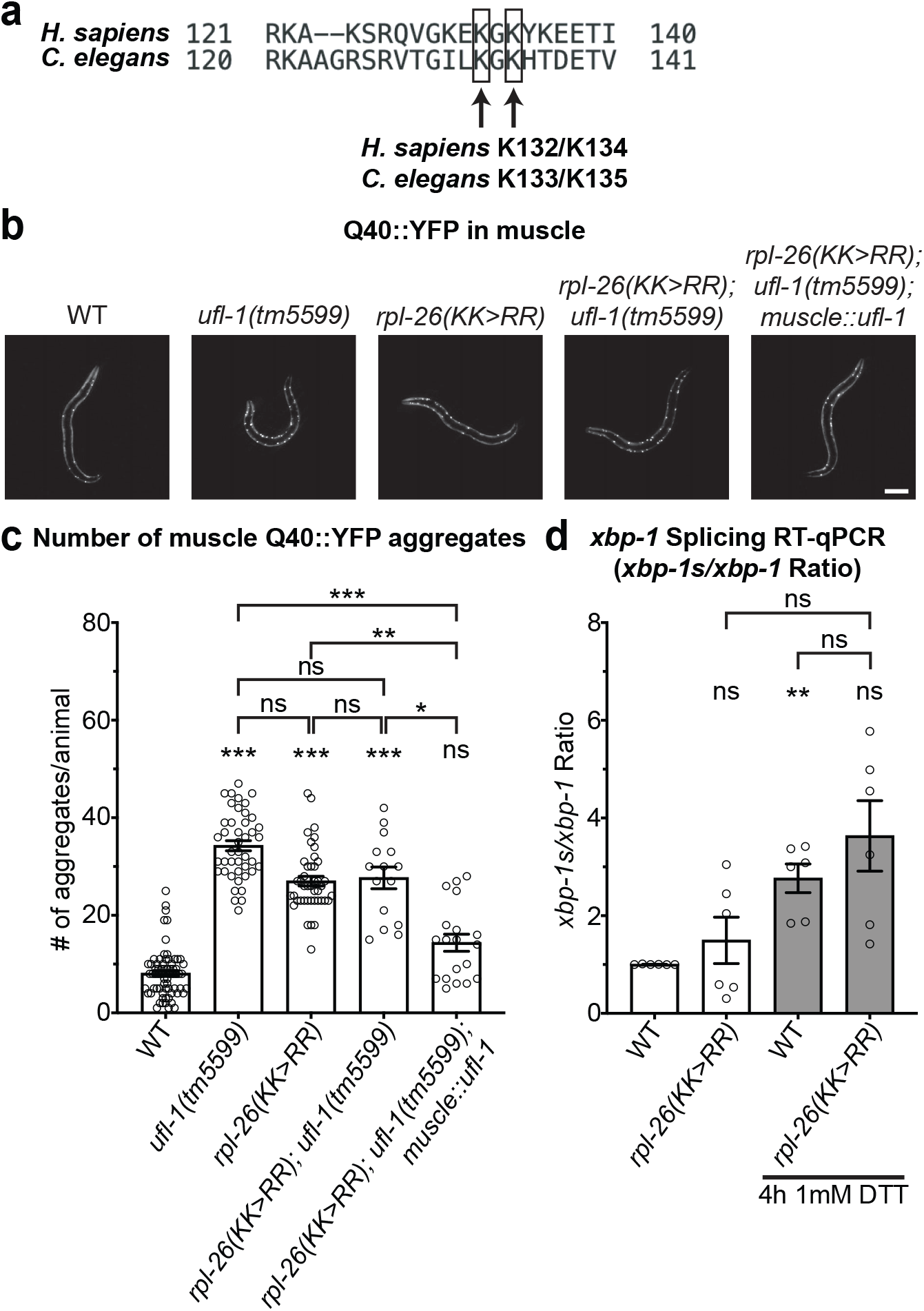
*rpl-26* mediates the increased Q40::YFP aggregates observed in UFM-1 pathway mutants. (a) Comparison of human and *C. elegans* UFMylation sites on RPL-26. K132/K134 in human RPL26 corresponds to K133/K135 in *C. elegans* RPL-26. (b) Representative images of Q40::YFP in muscle (*rmIs133*) in wild-type, *ufl-1(tm5599)*, *rpl-26(KK>RR)*, *rpl-26(KK>RR); ufl-1(tm5599)*, and *rpl-26(KK>RR); ufl-1(tm5599); muscle::ufl-1* animals. Images were enhanced for display purposes. Scale bar represents 100µm. (c) Quantification of the number of Q40::YFP aggregates per animal in age-matched L4 wild-type (Q40::YFP in muscle) (n=68), *ufl-1(tm5599)* (n=43), *rpl-26(KK>RR)* (n=43), *rpl-26(KK>RR); ufl-1(tm5599)* (n=15), and *rpl-26(KK>RR); ufl-1(tm5599); muscle::ufl-1* (n=19) animals. Kruskal-Wallis test with Dunn’s Multiple Comparisons Test revealed a significant difference between wildtype and *ufl-1(tm5599)* (p < 0.0001), wildtype and *rpl-26(KK>RR)* (p < 0.0001), wildtype and *rpl-26(KK>RR); ufl-1(tm5599)* (p < 0.0001), *ufl-1(tm5599)* and *rpl-26(KK>RR); ufl-1(tm5599); muscle::ufl-1* (p < 0.0001), *rpl-26(KK>RR)* and *rpl-26(KK>RR); ufl-1(tm5599); muscle::ufl-1* (p = 0.0043), and *rpl-26(KK>RR); ufl-1(tm5599)* and *rpl-26(KK>RR); ufl-1(tm5599); muscle::ufl-1* (p = 0.0191). (d) Ratio of relative expression of *xbp-1s* mRNA to total *xbp-1* mRNA (each normalized to *act-1*) under baseline and DTT-treated conditions (1mM DTT for 4h in M9 buffer) measured by RT-qPCR for wild-type and *rpl-26(KK>RR)* animals. n = 6 independent trials, each data point represents the average of 2-3 technical replicates within a trial. Brown-Forsythe and Welch ANOVA tests with Dunnett’s Multiple Comparisons Test revealed a significant difference between wildtype after DTT treatment compared to wildtype at baseline (p = 0.0087). Unless otherwise noted, all graphs show mean ± SEM. Differences versus wildtype are shown above each bar. All other comparisons are marked by brackets. ns, p > 0.05; *, p < 0.05; **, p < 0.01; ***, p < 0.001.

We next tested whether the effects of overexpressed *ufl-1* on Q40::YFP aggregates could be blocked by *rpl-26(KK>RR)*. We showed above that overexpression of wild-type *ufl-1* in muscle restores the number of Q40::YFP aggregates observed in *ufl-1* mutants back to wild-type levels (Fig. 3). If RPL-26 is the only target downstream of UFL-1 that regulates polyQ aggregates, then overexpressing wild-type *ufl-1* in muscle in *rpl-26(KK>RR); ufl-1(tm5599)* double mutants should have no effect on the number of Q40::YFP aggregates. In contrast, we found that expression of *ufl-1* in the muscle of *rpl-26(KK>RR); ufl-1(tm5599)* double mutants was sufficient to restore the number of muscle Q40::YFP aggregates back to wild-type levels (Fig. 4b and c, 1^st^ vs 5^th^ bars). This result suggests that overexpression of *ufl-1* can regulate Q40::YFP aggregates even in the absence of putative UFMylation sites in RPL-26. Taken together, our genetic double mutant data suggest that under basal conditions *rpl-26(KK>RR)* mediates the effects of *ufl-1* on Q40::YFP aggregates. However, our rescue data suggest that when *ufl-1* is overexpressed, or perhaps activated under physiological conditions, it is able to bypass RPL-26 and UFMylate one or more other substrates to regulate Q40::YFP aggregates.

Our data show that loss of the UFM-1 pathway results in activation of the UPR^ER^ under proteotoxic stress conditions (Fig. 1) and that loss of the UFM-1 pathway leads to increased Q40::YFP aggregates in a process that is mediated by *rpl-26* (Fig. 4). Thus, we tested whether RPL-26(KK>RR) mutants also enhance the UPR^ER^ under stress conditions. As described above, we monitored UPR^ER^ activation by measuring the levels of endogenous *xbp-1s* relative to total *xbp-1* transcript levels using RT-qPCR. Surprisingly, we found that *rpl-26(KK>RR)* mutants did not show a significant change in the ratio of *xbp-1s*/*xbp-1* under basal conditions or in the presence of DTT compared to their respective wild-type controls (Fig. 4d). These data suggest that, in contrast to *ufl-1* mutants treated with DTT (Fig. 1g), loss of the ability to UFMylate RPL-26 does not enhance the UPR^ER^ under stress conditions. Taken together, these data suggest that UFL-1-regulation of polyQ aggregates is mediated by RPL-26, but the ability of UFL-1 to regulate the UPR^ER^ is likely mediated by one or more other targets.

### *ufl-1* mutants have altered Q40::YFP aggregate composition

Our data show that loss of the UFM-1 pathway leads to an increase in the number of Q40::YFP aggregates and enhanced activation of the UPR^ER^ under stress conditions, consistent with altered protein homeostasis (Fig. 1 and Fig. 2). While enhanced activation of the UPR^ER^ under proteotoxic stress is expected to be beneficial for organismal proteostasis, it is not as clear whether the increased numbers of polyQ aggregates that we observe in UFM-1 pathway mutants is beneficial or detrimental to the animal. Some studies suggest that increased numbers of polyQ aggregates are detrimental to cellular and organismal health (Kazemi-Esfarjani and Benzer 2000; Morley et al. 2002; Sakahira et al. 2002; Nollen et al. 2004; Brignull et al. 2006; Kaiser et al. 2026). However, other studies suggest that increased numbers of aggregates may be beneficial for the organism due to the increased sequestration of polyQ species in aggregates that results in a decrease in soluble toxic oligomers in the cytoplasm (Arrasate et al. 2004; Kim et al. 2016; Van Pelt and Truttmann 2025; Bruck-Haimson et al. 2026). Additionally, aggregates contain a mixture of soluble and less toxic insoluble components and the proportion of these species within aggregates can impact their overall toxicity to the organism (Arrasate et al. 2004; Kim et al. 2016). Thus, we analyzed the composition of the Q40::YFP aggregates we observe in *ufl-1* mutants using Fluorescence Recovery After Photobleaching (FRAP). FRAP has previously been used to estimate the mobility of Q40::YFP aggregates in *C. elegans* (Morley et al. 2002; Brignull et al. 2006; Garcia et al. 2007; Beam et al. 2012; Moronetti Mazzeo et al. 2012). Briefly, half of a single Q40::YFP aggregate is photobleached and YFP fluorescence recovery is monitored every two minutes for a total of 10 minutes. The rate of YFP fluorescence recovery can be attributed to diffusion of mobile Q40::YFP from the unbleached side of the aggregate, providing an estimate of the fraction of mobile to immobile Q40::YFP in each aggregate, which may reflect differences in the soluble to insoluble proportion of the aggregate. We found that after photobleaching, Q40::YFP aggregates in wild-type animals recovered to ∼40% of pre-bleach levels following a 10-minute recovery period (Fig. 5a and b). In contrast, in *ufl-1* mutants, Q40::YFP aggregates recovered to ∼62% of pre-bleach levels (Fig. 5a and b). Taken together, our results indicate that loss of the UFM-1 pathway results in an increase in the number of polyQ aggregates and a change in their aggregation state with a higher proportion of soluble Q40::YFP per aggregate.

**Fig. 5.**
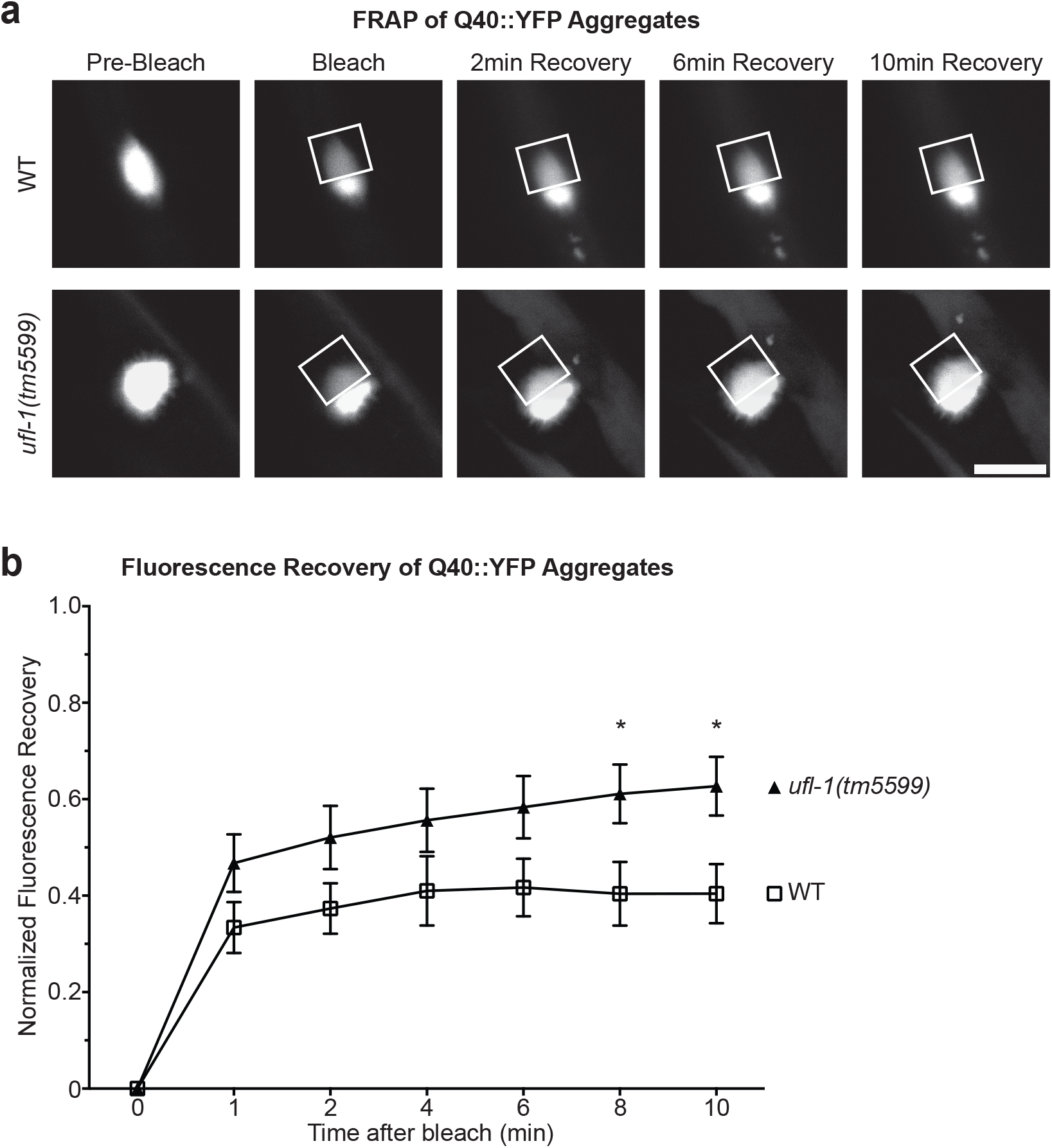
*ufl-1* mutants have altered polyQ aggregate composition. (a) Representative prebleach, bleach, 2-minute recovery, 6-minute recovery, and 10-minute recovery images of Q40::YFP in muscle (*rmIs133*) in age-matched L4 animals in wild-type (Q40::YFP in muscle) or *ufl-1(tm5599)* animals. Bleached region is boxed in white on Bleach and Recovery images. Scale bar represents 10µm. (b) Quantification of the normalized rate of recovery for muscle Q40::YFP aggregates (normalized to prebleach intensities) in wild-type (n=19) or *ufl-1(tm5599)* (n=20) animals. Two-way ANOVA with Tukey’s multiple comparisons test revealed a significant difference between genotypes at 8 minutes post-bleach (p = 0.0268) and 10 minutes post-bleach (p = 0.0141). Unless otherwise noted, all graphs show mean ± SEM. Differences between genotypes are shown above the corresponding time point. *, p < 0.05.

### *ufl-1* mutants exhibit enhanced intestinal healthspan and survival under stress

Given that loss of the UFM-1 pathway regulates proteostasis, we tested whether loss of *ufl-1* has beneficial or detrimental effects on organismal physiology using three distinct healthspan assays.

First, we measured the rate of contraction of the pharyngeal muscles (pharyngeal pumping rate) which is known to decline with age (Croll et al. 1977; Kenyon et al. 1993; Huang et al. 2004). We found that *ufl-1* mutants exhibit decreased pharyngeal pumping rates compared to wild-type animals at the L4 stage and adult day 11, but no difference at adult day 8 (Fig. 6a). We also tested whether expression of Q40::YFP in body wall muscle affected pharyngeal pumping as disruption of proteostasis in one tissue is known to affect other tissues (Taylor and Dillin 2013; Laranjeiro et al. 2019; Hoppe and Cohen 2020). We observed a decline in pharyngeal pumping rate in *ufl-1* mutants expressing muscle Q40::YFP compared to wild-type animals at the L4 stage. Interestingly, we observed an increase in pharyngeal pumping rate of *ufl-1* mutants expressing muscle Q40::YFP at adult day 8, although this increase was no longer apparent by adult day 11 (Supplementary Fig. 3a), perhaps because expression of Q40::YFP combined with increased age ultimately overwhelms the organism’s ability to handle proteotoxic stress. These data suggest that in wild-type animals, loss of *ufl-1* does not enhance pharyngeal healthspan and is actually detrimental in older adults. However, in the presence of muscle Q40::YFP, which causes proteotoxic stress itself, there appears to be a short window of time around adult day 8 where loss of *ufl-1* enhances pharyngeal function.

**Fig. 6.**
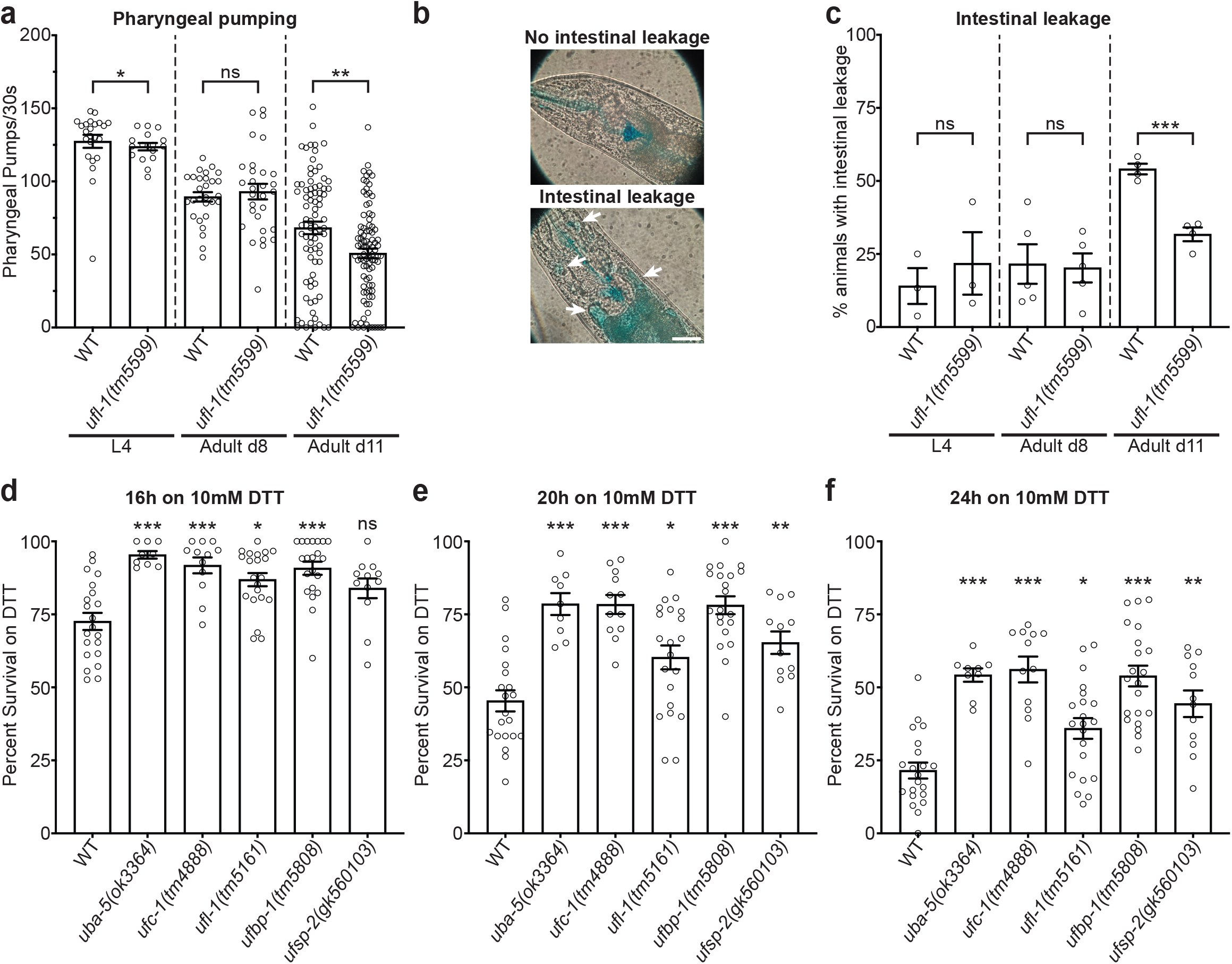
Loss of UFM-1 pathway genes results in enhanced healthspan and survival under stress. (a) Pharyngeal pumping rate of wild-type and *ufl-1(tm5599)* animals at L4, adult day 8 (d8), and adult day 11 (d11). n = 16-93 animals. Mann-Whitney test revealed a significant difference between wildtype and *ufl-1(tm5599)* at L4 (p = 0.0469). Unpaired t-test with Welch’s correction revealed a significant difference between wildtype and *ufl-1(tm5599)* at adult d11 (p = 0.0015) but not adult d8 (p = 0.5693). (b) Representative images of animals with an intact intestinal barrier (top) and intestinal barrier leakage (bottom). Arrows indicate intestinal leakage. Scale bars represent 100µm. (c) Percentage of animals with intestinal barrier leakage in wild-type and *ufl-1(tm5599)* animals at L4, adult day 8 (d8), and adult day 11 (d11). n = 3-5 independent trials. Unpaired t-test with Welch’s correction revealed a significant difference between wildtype and *ufl-1(tm5599)* at adult d11 (p = 0.0004) but not L4 (p = 0.5725) or adult d8 (p = 0.8773). (d-f) Percent survival of young adult wild-type, *uba-5(ok3364)*, *ufc-1(tm4888)*, *ufl-1(tm5161)*, *ufbp-1(tm5808)*, and *ufsp-2(gk560103)* animals after exposure to 10mM DTT for 16h (d), 20h (e), and 24h (f). n = 3-7 independent trials with three plates for each genotype. Each data point represents one plate of 15-20 animals. (d) Kruskal-Wallis test with Dunn’s multiple comparisons test revealed a significant difference between wildtype compared to: *uba-5(ok3364)* (p < 0.0001), *ufc-1(tm4888)* (p = 0.0003), *ufl-1(tm5161)* (p = 0.0112), and *ufbp-1(tm5808)* (p < 0.0001). (e) Brown-Forsythe and Welch ANOVA tests with Dunnett’s Multiple Comparisons Test revealed a significant difference between wildtype compared to: *uba-5(ok3364)* (p < 0.0001), *ufc-1(tm4888)* (p < 0.0001), *ufl-1(tm5161)* (p = 0.0467), *ufbp-1(tm5808)* (p < 0.0001), and *ufsp-2(gk560103)* (p = 0.0040) mutant animals. (f) Brown-Forsythe and Welch ANOVA tests with Dunnett’s Multiple Comparisons test revealed a significant difference between wildtype compared to: *uba-5(ok3364)* (p < 0.0001), *ufc-1(tm4888)* (p < 0.0001), *ufl-1(tm5161)* (p = 0.0130), *ufbp-1(tm5808)* (p < 0.0001), and *ufsp-2(gk560103)* (p = 0.0019)mutant animals. Unless otherwise noted, all graphs show mean ± SEM. Differences between genotypes at different ages are marked by brackets (a, c). Differences versus wildtype are shown above each bar (d-f). ns, p > 0.05; *, p < 0.05; **, p < 0.01; ***, p < 0.001.

Second, we analyzed intestinal barrier integrity using a dye permeability assay. The *C. elegans* intestine is a critical tissue for maintaining animal health and aging (Herndon et al. 2002; Libina et al. 2003; McGhee 2007; Gelino et al. 2016). Intestinal barrier integrity declines with age and can be used to monitor gut health in *C. elegans* and *D. melanogaster* (Herndon et al. 2002; Rera et al. 2012; Gelino et al. 2016; Laranjeiro et al. 2019). Briefly, the animals are fed a non-absorbable blue dye and the percentage of animals with dye leakage outside of the intestinal lumen into the body cavity is quantified (Fig. 6b). We found no change in the percentage of animals with intestinal dye leakage in *ufl-1* mutants compared to wildtype at the L4 stage or at adult day 8 (Fig. 6c). However, at adult day 11, ∼60% of wild-type animals exhibited dye leakage into the body cavity whereas only 35% of *ufl-1(tm5599)* mutants had dye leakage (Fig. 6c). This result suggests that loss of *ufl-1* results in improved intestinal barrier integrity in aged animals. We also tested if expression of Q40::YFP in muscle affected the intestinal barrier. We found that loss of *ufl-1* did not affect intestinal barrier integrity in polyQ-expressing animals at the L4 stage or adult day 8, but had a trending, albeit non-significant (p=0.06) decrease in dye-leakage at adult day 11 (Supplementary Fig. 3b). Together, these data suggest that in the absence of polyQ expression, loss of *ufl-1* enhances intestinal healthspan in adult day 11 aged animals.

Third, we performed an ER stress resistance assay, another important indicator of animal health, by monitoring survival in the presence of the protein-folding stressor DTT (Hertel et al. 2013; Park et al. 2017; Torres et al. 2022; Kaiser et al. 2026). We exposed young adult animals to 10mM DTT and assessed the percent survival after 16h, 20h, or 24h. Interestingly, we observed enhanced survival following DTT exposure in loss-of-function mutants of *uba-5* (E1 enzyme), *ufc-1* (E2 enzyme), *ufl-1* (E3 enzyme), and *ufbp-1* (ER anchored adaptor) at all time points (Fig. 6d-f). Loss of *ufsp-2* (cysteine protease) did not alter survival following exposure to DTT after 16h but did enhance survival after 20h and 24h (Fig. 6d-f). These data show that loss of UFM-1 pathway components results in increased survival under proteotoxic stress. This increase in survival in the presence of stress is likely due to increased activation of the UPR^ER^ in UFM-1 pathway mutants under stress (Fig. 1). Taken together, our data show that loss of UFM-1 pathway genes can be beneficial for the organism, enhancing intestinal healthspan in aged animals and increasing survival following exposure to ER stress.

### Loss of *ufl-1* extends lifespan in Q40::YFP animals

The healthspan enhancements we observed in UFM-1 pathway mutants led us to investigate whether loss of the UFM-1 pathway extended lifespan. We observed a moderate 9% increase in lifespan in *ufl-1* mutants (median lifespan: 24 days) compared to wild-type animals (median lifespan: 22 days) (Fig. 7 and Supplementary Table 3). Since expression of polyQ proteins increases proteotoxic stress and decreases lifespan (Morley et al. 2002; Gidalevitz et al. 2006; Nedosekin et al. 2021; Van Pelt and Truttmann 2025), we tested whether loss of *ufl-1* could suppress the proteotoxic effect of polyQ on lifespan. Consistent with prior studies (Nedosekin et al. 2021; Van Pelt and Truttmann 2025), we found that expression of Q40::YFP in muscle decreases lifespan (median lifespan: 15 days) compared to wild-type animals (median lifespan: 22 days) (Fig. 7 and Supplementary Table 3). Intriguingly, we found that the decrease in lifespan observed in Q40::YFP-expressing animals is suppressed in *ufl-1* mutants resulting in a ∼26% lifespan extension compared to Q40::YFP-expressing control animals, although their lifespan is still shortened compared to wild-type animals (Fig. 7, Supplementary Table 3, median lifespan: wild-type = 22 days, *Q40::YFP* animals = 15 days, *ufl-1; Q40::YFP* animals = 19 days). Thus, loss of the UFM-1 pathway extends lifespan and counteracts the reduction in lifespan caused by expression of expanded-polyQ proteins.

**Fig. 7.**
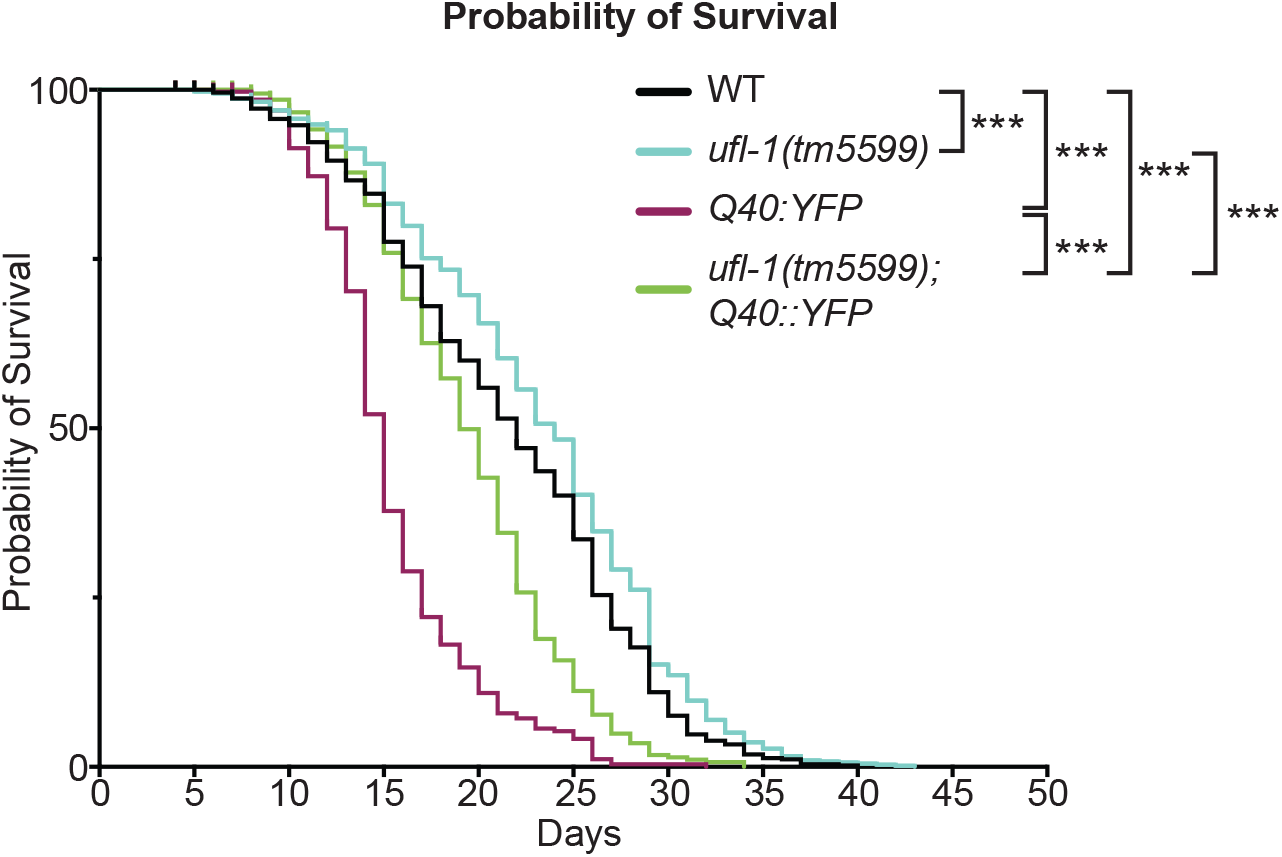
Loss of *ufl-1* extends lifespan. Kaplan-Meier survival curves for wild-type, *ufl-1(tm5599)*, Q40::YFP, and *ufl-1(tm5599); Q40::YFP* animals. Median lifespan: wildtype: 21 days, *ufl-1(tm5599)*: 21 days, *Q40::YFP*: 15 days, *ufl-1(tm5599); Q40::YFP*: 19 days. n=3-6 independent trials. Pairwise Mantel-Cox tests revealed significant differences between: wildtype and *Q40::YFP* (p < 0.0001); wildtype and *ufl-1(tm5599); Q40::YFP* (p < 0.0001); *Q40::YFP* and *ufl-1(tm5599); Q40::YFP* (p < 0.0001); and *ufl-1(tm5599)* and *ufl-1(tm5599); Q40::YFP* (p < 0.0001). Unless otherwise noted, all graphs show mean ± SEM. Differences are marked by brackets. ***, p < 0.0001.

## Discussion

In this study, we investigated the role of the UFM-1 pathway in regulating proteostasis and organismal health in *C. elegans* by analyzing loss-of-function mutants in individual components of the pathway. Consistent with prior studies (Hertel et al. 2013; Kaiser et al. 2026), we found that UFM-1 pathway mutants exhibit enhanced activation of the UPR^ER^ under conditions of proteotoxic stress (Fig. 1). Analysis of a polyQ aggregation model revealed that loss of UFM-1 pathway components resulted in an increase in the number of polyQ aggregates in muscle and a change in their composition (Fig. 2, Fig. 5), indicating that the UFM-1 pathway regulates protein homeostasis. Tissue-specific rescue experiments show that the UFM-1 E3 ligase *ufl-1* acts cell autonomously in muscle and non-cell autonomously in the nervous system to regulate polyQ aggregates in muscle (Fig. 3). We found that the increase in polyQ aggregates observed in *ufl-1* mutants is mediated by the large ribosomal subunit RLP-26 (Fig. 4), which is a major UFMylation target in mammalian cells (Walczak et al. 2019; Wang et al. 2020). Interestingly, loss of the UFM-1 pathway is beneficial at the organismal level as animals lacking UFM-1 pathway components exhibited improved intestinal barrier integrity with age, increased survival under stress (Fig. 6) and enhanced lifespan in polyQ-expressing animals (Fig. 7). Together, our findings suggest that loss of the UFM-1 pathway is beneficial for organismal health and aging under conditions of proteotoxic stress (i.e., chemical DTT treatment, aging or polyQ expression).

We monitored protein homeostasis in *C. elegans* using a well-established polyQ aggregation model that is just above the aggregation threshold (Q40::YFP)(Morley et al. 2002; Brignull et al. 2006). We found that mutants lacking each individual component of the core UFM-1 pathway results in increased numbers of Q40::YFP aggregates in muscle (Fig. 2), suggesting that loss of UFMylation regulates polyQ aggregates. Our results are consistent with two recent studies in *C. elegans* that used distinct polyglutamine aggregation models. After knockdown of *ufm-1* with RNAi, Bruck-Haimson et al. observed an increased number of polyQ aggregates based on a reporter using YFP-tagged with 35 glutamines (Q35::YFP) (Bruck-Haimson et al. 2026), and Kaiser et al. found an increase in aggregate size based on a reporter consisting of a fragment of Huntingtin tagged with YFP and 128 glutamines (Htt513(Q128)::YFP) (Kaiser et al. 2026). Our findings, together with these studies, show that loss of the UFM-1 pathway affects protein homeostasis and leads to an increase in protein aggregates.

### The UFM-1 pathway and RPL-26 act at the ER to regulate polyQ aggregates

Interestingly, we found that loss of the ER-anchored proteins, UFBP-1, which is part of the UFL-1 E3 ligase complex and essential for UFMylation, or ODR-4, which recruits UFSP-2 to the ER, is sufficient to increase the number of Q40::YFP aggregates (Fig. 2). These results suggest that the UFM-1 pathway acts at the ER to regulate polyQ aggregates in the cytoplasm. One of the most well characterized and major target of UFMylation in mammalian cells is the large ribosomal subunit protein RPL26 (Walczak et al. 2019; Wang et al. 2020; Scavone et al. 2023; Mihailovic et al. 2026). RPL26 is involved in ribosome-associated protein quality control at the ER and the selective degradation of damaged sections of the ER by autophagy, a process called ERphagy (Liang et al. 2020; Ishimura et al. 2022; Komatsu et al. 2024). UFMylation of RPL26 occurs when ribosomes stall due to faulty mRNA or defective nascent chains stuck in the translocon, and results in the removal of the nascent chain and disassembly of the ribosome from the translocon (Walczak et al. 2019; DaRosa et al. 2024; Makhlouf et al. 2024). We found that mutation of two conserved lysines (required for UFMylation of mammalian RPL26) to arginines in endogenous *rpl-26* (*rpl-26(KK>RR))* results in increased Q40::YFP aggregate number to a similar extent to that observed in *ufl-1* mutants (Fig. 4). These data suggest that loss of UFMylation of RPL-26 is sufficient to increase polyQ aggregates in muscle. Furthermore, the increase in aggregate number observed in *rpl-26(KK>RR); ufl-1* double mutants was not additive (Fig. 4). These data indicate that *rpl-26(KK>RR)* and *ufl-1* act in the same genetic pathway to regulate polyQ aggregates and are consistent with a model where UFL-1 acts at the ER to UFMylate RPL-26 to regulate Q40::YFP aggregates in the cytoplasm. Further studies will be required to determine whether the inability to UFMylate RPL-26 effects polyQ aggregates via an indirect mechanism by disrupting ERQC or a more direct mechanism as has been shown for FIC-1-mediated ampylation (Van Pelt and Truttmann 2025).

Our genetic double mutant data strongly suggest that RPL-26 is the major target downstream of UFL-1 that regulates polyQ aggregates. This finding is consistent with a mammalian study showing that RPL-26 is a major physiological UFMylation target in mammalian cells (Walczak et al. 2019). However, we also found that expression of wild-type *ufl-1* in *rpl-26(KK>RR); ufl-1* double mutants, was able to rescue the number of aggregates back to wild-type levels (Fig 4), suggesting that UFL-1 can regulate Q40::YFP aggregates even in the absence of RPL-26 as an UFMylation target. Taken together with our genetic double mutant analysis, our data suggest that under basal conditions, RPL-26 is the major target UFMylated by UFL-1 to regulate Q40::YFP aggregates. However, under some circumstances, such as when *ufl-1* is overexpressed or perhaps when the UFMylation machinery is upregulated or activated under physiological conditions, UFL-1 may be able to bypass RPL-26 and UFMylate one or more other targets to regulate Q40::YFP aggregates.

In addition to regulating polyQ aggregates, our data show that loss of the UFM-1 pathway leads to increased UPR^ER^ activation under conditions of proteotoxic stress (i.e, after treatment with DTT)(Fig. 1g). In contrast, we found that *rpl-26(KK>RR)* mutants did not alter the UPR^ER^ (Fig. 4d) under basal conditions or in the presence of DTT based on RT-qPCR measurements of *xbp-1s*, suggesting that there are likely other substrates downstream of *ufl-1* that mediate the effects of the UFM-1 pathway on the UPR^ER^. These other substrates may enhance the UPR^ER^ by upregulating the expression of UPR^ER^ components (Zhang et al. 2012; Hertel et al. 2013; Cai et al. 2015; Kaiser et al. 2026). Additional targets may mediate the effects of the UFM-1 pathway on other aspects of the PN. For example, a recent study identified additional UFMylated substrates that also regulate proteostasis including nucleolar FIB-1 and NOL-56 in ASI neurons and CAR-1 and CGH-1 in the germline (Bruck-Haimson et al. 2026).

### The UFM-1 pathway acts cell-autonomously and non-cell autonomously to polyQ aggregates

Prior studies have shown that organismal proteostasis is regulated by inter-tissue signals (Taylor and Dillin 2013; Frakes et al. 2020; Özbey et al. 2020; Bruck-Haimson et al. 2026). Consistent with those studies, our results show that *ufl-1* can regulate polyQ aggregates in muscle both cell autonomously and non-cell autonomously (Fig. 3). The UFM-1 pathway enzymes were previously reported to be predominantly expressed in the intestine (Hertel et al. 2013; Kaiser et al. 2026). However, we found that expression of wild-type *ufl-1* specifically in the intestine of *ufl-1* mutants did not alter the numbers of Q40::YFP aggregates in muscle (Fig 3). Although, it will be informative to test if re-expression of wild-type *ufl-1* in the intestine eliminates the beneficial effects of *ufl-1* mutation on intestinal barrier integrity. Single-cell RNAseq studies indicate that UFM-1 pathway genes are widely expressed in many other tissues including the nervous system and muscle (Hammarlund et al. 2018; Taylor et al. 2021). We found that re-expression of wild-type *ufl-1* in the muscle of *ufl-1* mutants reverses aggregate number back to baseline suggesting that the UFM-1 pathway regulates muscle Q40::YFP aggregates cell autonomously (Fig. 3). Interestingly, re-expression of wild-type *ufl-1* in neurons of *ufl-1* mutants completely reverses aggregate number back to baseline (Fig. 3), suggesting that the UFM-1 pathway can also act in neurons to regulate polyQ aggregates in muscle. These data reveal an intriguing inter-tissue signal between neurons and muscle. It will be interesting in the future to test whether this inter-tissue signal is mediated by tyramine as has been shown for proteostatic signals between neurons and the intestine (Özbey et al. 2020) or via some other mechanism. Together, these data are consistent with the UFM-1 pathway functioning cell-autonomously and non-cell autonomously in multiple tissues to regulate proteostasis.

### Effects of the UFM-1 pathway on polyQ aggregate composition, healthspan and lifespan

Our results show that loss of UFM-1 pathway components leads to an increase in the number of polyQ aggregates. However, it is not clear if this increase in aggregates is beneficial or detrimental for cellular or organismal physiology and health. Canonical aggregation-toxicity models suggest that increased aggregate burden reflects impaired proteostasis, increased toxicity and functional decline (Kazemi-Esfarjani and Benzer 2000; Morley et al. 2002; Sakahira et al. 2002; Nollen et al. 2004; Brignull et al. 2006; Kaiser et al. 2026). Consistent with this, some studies suggest that reduced UFMylation results in impaired motility and increased aggregates leading to detrimental physiological consequences in flies and worms such as shortened lifespan (Duan et al. 2016; Kaiser et al. 2026). In contrast, other studies suggest that increased numbers of aggregates represents increased sequestration of potentially toxic aggregation prone species and is thus beneficial for organismal health and lifespan (Arrasate et al. 2004; Kim et al. 2016; Van Pelt and Truttmann 2025; Bruck-Haimson et al. 2026). For example, Arrasate et al. showed that increased sequestration of Huntingtin (Htt) protein in inclusion bodies decreases levels of diffuse, toxic mutant Htt protein in the cytoplasm and protects neurons from death (Arrasate et al. 2004). Consistent with this, another study showed that soluble oligomeric polyQ-expanded Htt species, generated from polyQ monomers, correlate with ER stress and are more toxic than large Htt aggregates (Leitman et al. 2013). Leitman et al. suggested a model where sequestration of polyQ monomers within large aggregates is beneficial because the monomers are prevented from forming toxic oligomers in the cytoplasm. Additionally, proteomic studies showed that soluble oligomeric Htt species are more toxic than insoluble Htt species found in large inclusions because the soluble oligomers interact with and disrupt more key proteins involved in critical cellular pathways (Arrasate et al. 2004; Kim et al. 2016). Together, these studies suggest that increased aggregates may be beneficial because it results in the sequestration of polyQ species in aggregates that ultimately prevents the formation of toxic oligomers in the cytoplasm.

Our results are consistent with the idea that increased aggregates are beneficial for the organism. We found increased numbers of aggregates in *ufl-1* mutants and this correlated with increased healthspan and lifespan. In addition, our FRAP experiments showed that aggregates in *ufl-1* mutants are composed of a larger mobile fraction of Q40::YFP (Fig. 5), which is consistent with an increased proportion of soluble Q40::YFP in the aggregates. Taken together with other studies, our results suggest that increased aggregates in *ufl-1* mutants may be beneficial for the organism because it results in the increased sequestration or containment of soluble polyQ species in aggregates, perhaps preventing the formation of soluble more toxic aggregates in the cytoplasm. Our results are in line with a recent study showing that RNAi knockdown of *ufm-1* results in an increase in the number of Q35::YFP aggregates in muscle and these aggregates have altered composition (Bruck-Haimson et al. 2026). Additionally, they showed that RNAi knockdown of *ufm-1* suppresses the decrease in locomotion caused by expression of Q35::YFP in neurons, suggesting that loss of UFMylation is beneficial (Bruck-Haimson et al. 2026). In support of this, we observed enhanced healthspan parameters in UFM-1 pathway mutants. We found that UFM-1 pathway mutants exhibit enhanced activation of the UPR^ER^ under stress (Fig. 1), enhanced survival under ER stress (Fig. 6), and improved intestinal barrier integrity with age (Fig. 6). We also found that *ufl-1* mutants exhibit a modest increase in lifespan (Fig. 7). However, the beneficial effect of *ufl-1* mutation on lifespan was most dramatic in polyQ expressing animals, where *ufl-1* mutants suppressed the detrimental effects of polyQ expression on lifespan by 26% percent (Fig. 7). Although we acknowledge that we cannot directly connect the effects of UFM-1 pathway mutants on polyQ aggregates to the beneficial effects on healthspan and lifespan, our data are consistent with a model where increased sequestration of soluble polyQ species in aggregates is beneficial for the organism. Alternatively, because the UFM-1 pathway genes are broadly expressed in many tissues based on single cell RNAseq data (Hammarlund et al. 2018; Taylor et al. 2021) and our results show that the UFM-1 pathway has inter-tissue effects, it remains possible that the effects of *ufl-1* mutants on polyQ aggregates in muscle are uncoupled from the beneficial effects on healthspan and lifespan. In this latter scenario, loss of the UFM-1 pathway might enhance the UPR^ER^ in other non-muscle tissues leading to the beneficial effects on healthspan and lifespan.

## Concluding Remarks

UFM-1 pathway mutants exhibit enhanced activation of the UPR^ER^ in the presence of various forms of proteotoxic stresses and this may be beneficial for several measures of healthspan as well as lifespan. We propose that reduction of UFMylation causes a mild proteostatic stress that “primes” or activates other adaptive stress responses leading to protective proteostatic pathways that enhance organismal health and extend lifespan. This form of adaptive homeostasis is reminiscent of hormesis, which is a response that occurs when a low dose stress triggers an adaptive response that benefits the organism (Cypser et al. 2006; Kozlowski et al. 2014; Matai et al. 2019). Indeed, prior studies in *C. elegans* have shown that constitutive activation of the UPR^ER^ in neurons or glial cells, by expressing XBP-1s in a cell-specific manner, results in ER hormesis and inter-tissue signals that enhance ER stress resistance and lifespan (Taylor and Dillin 2013; Frakes et al. 2020). While all studies suggest that UFMylation is important for regulating proteostasis and aging, it remains unclear why certain perturbations of UFMylation promote protective responses while others cause detrimental effects (Zhang et al. 2012; Hertel et al. 2013; Cai et al. 2015; Duan et al. 2016; Bruck-Haimson et al. 2026; Kaiser et al. 2026). Our findings support a model in which the UFM-1 pathway acts at the ER to suppress the UPR^ER^ and regulate proteostasis and organismal health. The effect of the UFM-1 pathway on polyQ aggregates are mediated by RPL-26 whereas its effects on the UPR^ER^ are likely mediated by one or more other targets. Overall, the UFM-1 pathway appears to be detrimental for organismal health resulting in decreased survival under proteotoxic stress, decreased intestinal barrier integrity with age and decreased lifespan especially in the presence of chronic proteotoxic stress induced by expression of expanded polyQ proteins. Although it is clear that systemic inhibition of UFMylation is embryonic lethal in mice due to defects in hematopoiesis (Cai et al. 2015; Wei and Xu 2016) and can lead to neurodevelopmental diseases in humans (Colin et al. 2016; Muona et al. 2016; Nahorski et al. 2018), our findings suggest that selective inhibition of UFMylation in specific tissues after development is complete may enhance protein homeostasis with potential beneficial effects on healthspan and lifespan.

## Acknowledgements

We thank Rebecca Taylor and Thorsten Hoppe for strains and reagents. We thank Jon Madison for critical input and discussions during the early stages of this project. We thank Annette McGehee, Julia Yelick and Kaitlin Markoja for assistance generating the ufl-1 transgene. We thank Malavika Raman, Molly Hodul, Regina Powers and Elaina Cote for helpful discussions and critical comments on this manuscript. We thank Victor Hatini for assistance with the LSM800 confocal microscope. The Scientific color map “Hawaii” (Crameri 2018a) is used in this study (Crameri et al. 2020).

## Study Funding

This study was funded in part by a National Science Foundation grant IOS#2439125 (to P.J.) and the Dr. Marie Rozan Class of 1964 Gift for Neurodegeneration Research (to P.J.). Some strains were provided by the *Caenorhabditis* Genetics Center (CGC), which is funded by NIH Office of Research Infrastructure Programs (P40 OD010440), and by the National BioResource Project (NBRP) which is funded by the Japanese government.

## Data Availability Statement

Strains and reagents generated in this study are available upon request from the corresponding author P.J.

## Notes

### Competing Interest Statement

The authors have declared no competing interest.

