## Supplementary Figures and Tables for "Loss of the UFM-1 pathway enhances the ER Unfolded Protein Response under stress to improve proteostasis and organismal health in *Caenorhabditis elegans*"

### Supplementary Figure Legends

**Supplementary Fig. 1. The UFM-1 pathway and mutant allele information.**
(a) Schematic of the UFM-1 pathway. Like ubiquitination, addition of UFM-1 to lysine residues on target substrates (UFMylation) requires an E1-E2-E3 enzyme cascade. UBA-5 (E1 enzyme) will activate and bind UFM-1 and transfer it to the E2 enzyme, UFC-1. UFL-1 (E3 enzyme) acts together with the ER-anchored UFBP-1 and an adaptor such as CDK5-RAP3 (not shown) to recruit the target substrate and form a complex with UBA-5-UFC-1-UFM-1 to transfer UFM-1 onto the target substrate. This pathway also utilizes a cysteine protease, UFSP-2, that removes UFM-1 from the substrate, and two ER anchored adaptor proteins: UFBP-1, which is necessary for UFL-1 function, and ODR-4, which recruits UFSP-2 to the ER membrane to carry out its function.
(b) Exon-intron diagrams for UFM-1 pathway genes illustrating the genetic lesions for the various mutant alleles used in this study. Scale bars represent 100 base pairs.

**Supplementary Fig. 2. UFM-1 pathway genes do not regulate the *unc-54*/myosin promoter.**
(a) Representative images of diffuse Q24::YFP (*rmIs130*) in muscle of L4 wild-type (Q24::YFP in muscle), *uba-5(ok3364)*, and *ufl-1(tm5161)* animals. Scale bar represents 100μm.
(b) Quantification of normalized fluorescence intensity of Q24::YFP in muscle of L4 wild-type (n=54), *uba-5(ok3364)* (n=31), and *ufl-1(tm5161)* (n=35) animals. Kruskal-Wallis test with Dunn's Multiple Comparisons test revealed no significant difference between genotypes compared to wildtype (*uba-5(ok3364)*, p > 0.9999; *ufl-1(tm5161)*, p = 0.3642). Unless otherwise noted, all graphs show mean ± SEM. Differences versus wildtype are shown above each bar. ns, p > 0.05.

**Supplementary Fig. 3. Loss of *ufl-1* enhances pharyngeal healthspan in Q40::YFP-expressing animals**
(a) Pharyngeal pumping rate of animals expressing Q40::YFP in muscle (*rmIs133*) in wild-type and *ufl-1(tm5599)* animals at L4, adult day 8 (d8), and adult day 11 (d11). n = 14-82 animals. Mann-Whitney test revealed a significant difference between *Q40::YFP* and *ufl-1(tm5599); Q40::YFP* at L4 (p = 0.0037) but not adult d11 (p = 0.3784). Unpaired *t*-test with Welch's correction revealed a significant difference between *Q40::YFP* and *ufl-1(tm5599); Q40::YFP* at adult d8 (p = 0.0256).
(b) Percentage of animals with intestinal barrier leakage in animals expressing Q40::YFP in muscle (*rmIs133*) in wildtype and *ufl-1(tm5599)* animals at L4, adult day 8 (d8), and adult day 11 (d11). n = 6-7 independent trials. Unpaired *t*-test with Welch's correction revealed no significant difference between *Q40::YFP* and *ufl-1(tm5599); Q40::YFP* at L4 (p = 0.7877), adult d8 (p = 0.1627), or adult d11 (p = 0.0646).
Unless otherwise noted, all graphs show mean ± SEM. Differences between genotypes at different ages are marked by brackets. ns, p > 0.05; *, p < 0.05; **, p < 0.01.

Supplementary Figure 1

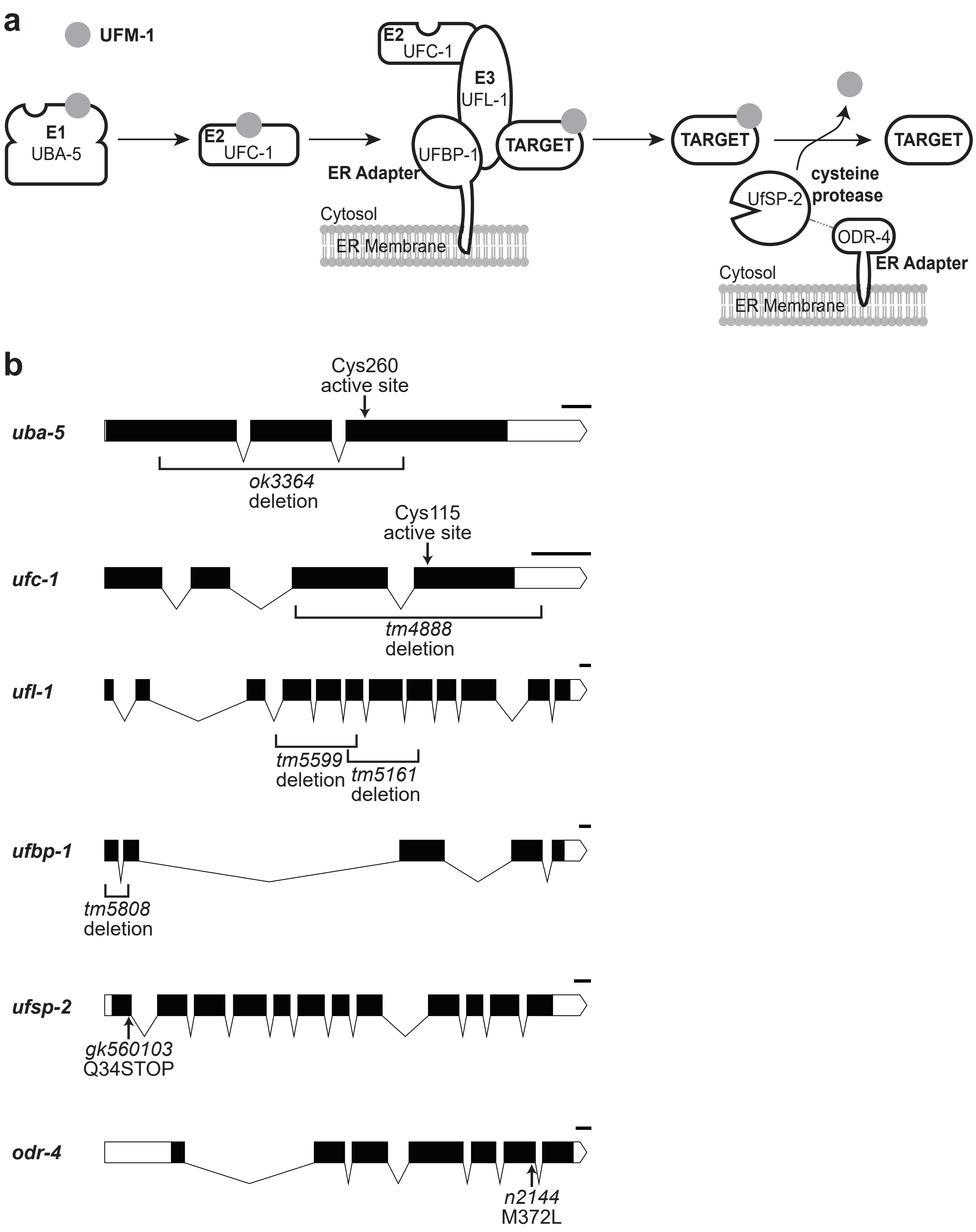

Supplementary Figure 2

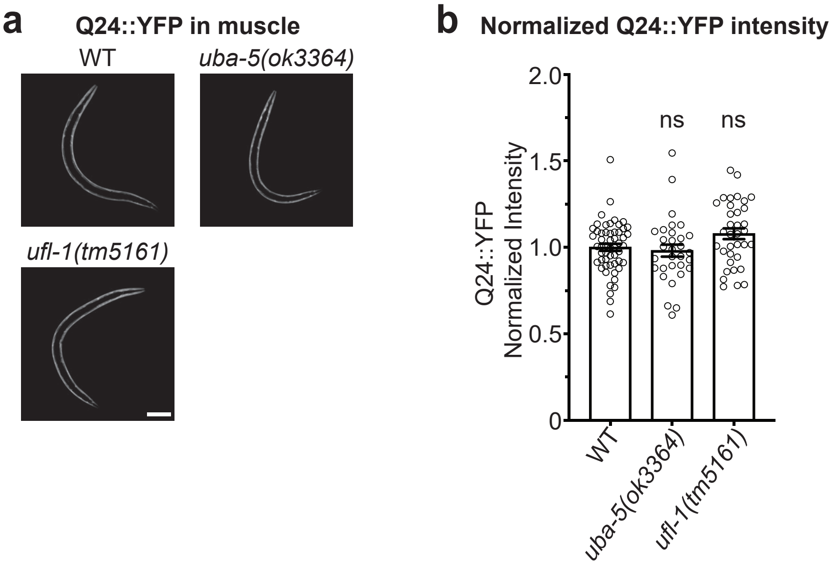

Supplementary Figure 3

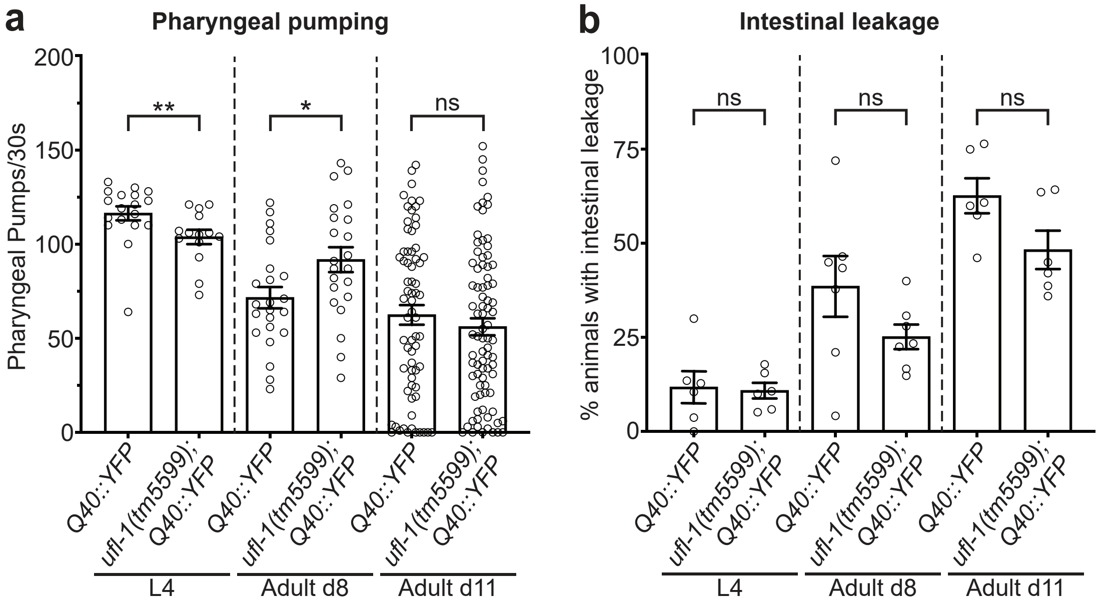

**Supplementary Table 1: Strains used in this study**

| Strain | Genotype | Source |
|---|---|---|
| N2 | Wildtype, Bristol | CGC |
| FJ1185 | *uba-5(ok3364)*6 I* | This study (CGC strain backcrossed to N2) |
| FJ2123 | *ufc-1(tm4888)*6 III* | This study (NBRP strain backcrossed to N2) |
| FJ1764 | *ufl-1(tm5161)*6 IV* | This study (NBRP strain backcrossed to N2) |
| FJ1454 | *ufl-1(tm5599)*3 IV* | This study (NBRP strain backcrossed to N2) |
| FJ1187 | *ufsp-2(gk560103)*6 IV* | This study (CGC strain backcrossed to N2) |
| FX05808 | *ufbp-1(tm5808) III* | NBRP |
| MT5300 | *odr-4(n2144) III* | CGC |
| AM141 | *rmIs133 [unc-54p::Q40::YFP] X* | CGC |
| FJ1747 | *uba-5(ok3364)*6 I; rmIs133 X* | This study |
| FJ2114 | *ufc-1(tm4888)*7 III; rmIs133 X* | This study |
| FJ1703 | *ufl-1(tm5161)*6 IV; rmIs133 X;* | This study |
| FJ1847 | *ufl-1(tm5599)*4 IV; rmIs133 X* | This study |
| FJ1758 | *ufsp-2(gk560103)*7 IV; rmIs133 X* | This study |
| FJ1815 | *ufbp-1(tm5808) III; rmIs133 X* | This study |
| FJ1816 | *odr-4(n2144) III; rmIs133 X* | This study |
| AM138 | *rmIs130 [unc-54p::Q24::YFP] II* | CGC |
| FJ2069 | *uba-5(ok3364) I; rmIs130 II* | This study |
| FJ1957 | *rmIs130 II; ufl-1(tm5161) IV* | This study |
| FJ2152 | *ufl-1(tm5599) IV; rmIs133 X; pzEx496 [myo-3p::ufl-1;unc-122p::GFP]* | This study |
| FJ2153 | *ufl-1(tm5599) IV; rmIs133 X; pzEx504 [vha-6p::ufl-1;unc-122p::GFP]* | This study |
| FJ2160 | *ufl-1(tm5599) IV; rmIs133 X; pzEx505 [snb-1p::ufl-1;unc-122p::GFP]* | This study |
| PP1386 | *hhIs113 [nhx-2p::cpl-1* $^{W32AW35A}$ *::YFP;myo-2p::mCherry] I (CPL-1*::YFP)* | Gift from Thorsten Hoppe (Finger et al. 2019) |
| FJ2117 | *hhIs113 I; ufl-1(tm5161)*7 IV* | This study |
| FJ2131 | *hhIs113; sel-11(tm1743)*7 V* | This study |
| RCT21 | *rmsIs8 [xbp-1p::xbp-1s::GFP;rol-6(su1006)]* | Gift from Rebecca Taylor (Özbey et al. 2020) |
| FJ2121 | *uba-5(tm5599)*7 I; rmsIs8* | This study |
| FJ2120 | *ufl-1(tm5599)*4 IV; rmsIs8* | This study |
| FJ2142 | *bpIs151 [sqst-1p::sqst-1::GFP;unc-76(+)]* | This study (CGC strain HZ589 outcrossed to N2) |
| FJ2115 | *uba-5(ok3364)*7 I; bpIs151* | This study |
| FJ2136 | *atg-18(gk378)*8; bpIs151* | This study |
| FJ2132 | *rpl-26(syb11091)*4 II (Also referred to as rpl-26(KK>RR))* | This study (Suny Biotech strain backcrossed to N2) |
| FJ2130 | *rpl-26(syb11091) II; rmIs133 X* | This study |
| FJ2151 | *rpl-26(syb11091) II; ufl-1(tm5599) IV; rmIs133 X* | This study |
| FJ2161 | *rpl-26(syb11091) II; ufl-1(tm5599) IV; rmIs133 X; pzEx496 [myo-3p::ufl-1;unc-122p::GFP]* | This study |

**Supplementary Table 2: Oligos used in this study**

**GENOTYPING**

| Primer Name | Strain | Sequence (5' > 3') | Purpose |
|---|---|---|---|
| uba-5_FWD | uba-5(ok3364) | AGT GTG ATG GAA GGC GAG AG | PCR genotyping |
| uba-5_REV | uba-5(ok3364) | TTT AAA CCG CCT TGG AAA TG | |
| ufc-1_FWD | ufc-1(tm4888) | CGT ACC GTC ACT TGG TGT AA | PCR genotyping |
| ufc-1_REV | ufc-1(tm4888) | GGG TAT CAA TGA CTG TCA CA | |
| ufl-1_FWD | ufl-1(tm5161) and ufl-1(tm5599 | GGC GCG TTG CTC TTT GTG TA | PCR genotyping |
| ufl-1_REV | ufl-1(tm5161) and ufl-1(tm5599 | CGC TGG CAA CCA ATT GAT GT | |
| ufbp-1_FWD | ufbp-1(tm5808) | GCA CCT TCA TTC ATT GCC AG | PCR genotyping |
| ufbp-1_REV | ufbp-1(tm5808) | GCG TCT TCC TCG CAT ACT AG | |
| ufsp-2_FWD | ufsp-2(gk560103) | ACC CGC TTC TTC CGT TTA TC | PCR genotyping and sequencing |
| ufsp-2_REV | ufsp-2(gk560103) | AGC GTA GGCGAA TAG CAC AT | |
| odr-4_FWD | odr-4(n2144) | CGC GCA GCT GTT CCA ATC AGA TC | PCR genotyping and sequencing |
| odr-4_REV | odr-4(n2144) | GCC AAG AGT CCG AT ATC AAT GC | |
| sel-11_FWD | sel-11(tm1743) | GGT AGA ACT CCG CCT GTG AT | PCR genotyping |
| sel-11_REV | sel-11(tm1743) | GTA AGA TGG GTT GAC TGG TG | |
| atg-18_FWD | atg-18(gk378) | CGA TGG TAG GCC TAA GTA GC | PCR genotyping |
| atg-18_REV | atg-18(gk378) | GAT TGA ACT TTA GAC ACG CG | |
| rpl-26_FWD | rpl-26(syb11091) | CAT AAG GGA AAC ACT GGA C | PCR genotyping and sequencing |
| rpl-26_REV | rpl-26(syb11091) | AAT TCC ACG CGT CTA CGT C | |

**SUBCLONING**

| Primer Name | Sequence (5' > 3') | Purpose |
|---|---|---|
| ufl-1 forward | GAT CGC TAG CAA AAA TGA CTT CCT GGG CAG ATA TT | Subcloning, primer contains NheI restriction site, used for FJ#155 and FJ#156 |
| ufl-1 reverse | GAT CGG TAC CTT ATG GCA TAA TTG ACT GTT TCA GA[illegible] | Subcloning, primer contains KpnI restriction site, used for FJ#155 and FJ#156 |
| ufl-1WTFP | GAT CGC TAG CAA AAA TGG CGG CCG CGA TGA CTT C[illegible] | Subcloning, primer contains NheI restriction site, used for FJ#154 |
| ufl-1WTRP | GAT CGG TAC CTT ATG GCA TAA TTG ACT GTT TCA GA[illegible] | Subcloning, primer contains KpnI restriction site, used for FJ#154 |
| ufl-1 internal 2 forward | GAA TGC CAG ACA ATT GGT TC | internal sequencing primer |
| ufl-1 internal 3 forward | CAT TTG GCT CTT CAA TCT C | internal sequencing primer |
| LD2 | GTT CTA TGT TAT GTT AGT ATC ATT CG | sequencing |
| LD3 | CGT ATG TTG TGT GGA ATT GTG AGC GG | sequencing |
| LD4 | GGT GGG AGC ACA GAC AAA GAG CAT GT | sequencing |
| LD5 | GGA CCC AAA GGT ATG TTT CGA ATG | sequencing |

**RT-qPCR**

| Primer Name | Gene | Sequence (5' > 3') | |
|---|---|---|---|
| xbp-1Tot_FWD | xbp-1 total | CAC CTC CAT CAA CAA CAA CAT | RT-qPCR |
| xbp-1Tot_REV | xbp-1 total | AAC CGT CTG CTC CTT CCT CAA | RT-qPCR |
| xbp-1s_FWD | xbp-1s | CGT GCC TTT GAA TCA GCA GTG | RT-qPCR |
| xbp-1s_REV | xbp-1s | CGA GGT GTC CAT CTT CTT GTT | RT-qPCR |
| act-1_FWD | act-1 | CCA GGA ATT GCT GAT CGT ATG CAG AA | RT-qPCR |
| act-1_REV | act-1 | TGG AGA GGG AAG CGA GGA TAG A | RT-qPCR |

| Supplementary Table 3: Lifespan Data | | | |
|---|---|---|---|
| **Strain** | **# Included** | **Median Lifespan (Lower 95% CI – Upper 95% CI)** | **% Life Span Change (Strains compared)** |
| Wildtype (N2) | 590/734 | 22 days (21 - 23 days) | - |
| *ufl-1(tm5599)* | 677/744 | 24 days (23 – 25 days) | 9.1% (vs wildtype) |
| *rmIs133* [Q40::YFP] | 281/359 | 15 days (14 – 15 days) | -31.8% (vs wildtype) |
| *ufl-1(tm5599); rmIs133* [Q40::YFP] | 304/362 | 19 days (19 – 20 days) | -13.6% (vs wildtype) |
| | | | 26.7% (vs rmIs133 [Q40::YFP]) |
